# Direct digital sensing of protein biomarkers in solution

**DOI:** 10.1101/2020.05.24.113498

**Authors:** Georg Krainer, Kadi L. Saar, William E. Arter, Timothy J. Welsh, Magdalena A. Czekalska, Raphaël P.B. Jacquat, Quentin Peter, Walther C. Traberg, Arvind Pujari, Akhila K. Jayaram, Pavankumar Challa, Christopher G. Taylor, Lize-Mari van der Linden, Titus Franzmann, Roisin M. Owens, Simon Alberti, David Klenerman, Tuomas P.J. Knowles

## Abstract

The detection of proteins is of central importance to biomolecular analysis and diagnostics, yet fundamental limitations due to the surface-based nature of most sensing approaches persist, and limited improvements have been designed to integrate multimodal information beyond concentration measurements. Here we present a single-molecule microfluidic sensing platform for digital protein biomarker detection in solution, termed digital immunosensor assay (DigitISA). DigitISA is based on microchip electrophoretic separation combined with single-molecule detection and enables absolute number–concentrations quantification of proteins in a single, solution-phase step. Applying DigitISA to a range of targets including amyloid aggregates, exosomes, and biomolecular condensates, we demonstrate that the assay provides information beyond stoichiometric interactions, and enables characterization of immunochemistry, binding affinity, and protein biomarker abundance. Together, DigitISA constitutes a new experimental paradigm for the digital sensing of protein biomarkers, and enables analyses of targets that would otherwise be hard or impossible to address by conventional immuno-sensing techniques.

## Introduction

The sensitive detection and quantitation of target biomolecules is essential in many areas of fundamental and applied science, ranging from biomolecular analysis and biophysics to clinical diagnostics. Proteins, in particular, are an important class of biomolecular targets, as they are ubiquitously regulated and affected in cellular physiology and disease, and as such can be used as biomarkers to diagnose and monitor pathological conditions as well as the efficacy of treatments. Developments in protein sensing are therefore a promising route toward more accurate protein analysis and disease diagnostics, with great potential in the emerging field of precision and personalized medicine.

Compared to other biomarker species, such as nucleic acid analytes, proteins are significantly more challenging to detect, as they cannot be amplified directly or targeted according to base-pair complementarity.^1, 2^ Instead, to achieve specificity and sensitivity, protein detection assays typically operate via surface-capture of target molecules by affinity reagents, which isolate the target protein prior to detection.^3, 4^ Foremost amongst sensing techniques that employ this principle are enzyme-linked immunosorbent assays (ELISAs),^5, 6^ which rely, in their most common implementation, on the surface-capture of target molecules by a dual antibody pair in a “sandwich” complex format, followed by an enzyme-driven signal amplification step. A number of recent approaches have advanced the classical ELISA technique, enabling remarkable improvements in its sensitivity and throughput.^7–10^ This has culminated in the development of digital sensing formats such as bead-based digital ELISAs developed by Quanterix,^11^ and methods such as those pioneered by Luminex,^12^ NanoString,^13^ and SomaSCAN,^14^ which display remarkable parallelization capabilities by simultaneously sensing a wide variety of protein targets down to attomolar sensitivities. Apart from ELISA and bead-based assays, other sensing techniques, including single-molecule and microfluidics-based assays, have found use in the quantitation and detection of proteins. These include methods such as fluorescence correlation spectroscopy, fluorescence anisotropy, microscale thermophoresis, surface-plasmon resonance and gel-based band-shift assays.^15–18^ Such methods, while lower in throughput, give the possibility to probe physical parameters of the interaction, and as such go beyond simple concentration measurements as is the case in classical ELISA or bead-based assays.

Despite the wide array of techniques available for protein sensing and the remarkable advances, in particular in terms of sensitivity, made over the past few decades, sensing techniques still bare significant and fundamental limitations. A central challenge common to ELISA and bead-based sensing techniques as well as other surface-based assay (e.g., surface plasmon resonance) is that these approaches remain reliant on surface-immobilization of target analyte molecules via capture probes. Surface-immobilization is problematic as it limits the analyte capture efficiency due to the finite binding area of the surface and thus makes the use of highly optimized capture probes, such as antibodies, with sub-nanomolar affinities necessary, whose production is often non-trivial and not feasible for some biomarker targets. In addition, surface- and matrix-based assays present challenges due to non-specificity binding, which makes these assays prone to false-positive signals. Even more significantly, surface- or matrix-based assays may not be appropriate for many analytes of diagnostic interest. Two examples of emerging interest include exosomes and biomolecular condensates. Their interaction with a surface or matrix (e.g., in ELISAs, bead-based or gel shift assays) may damage the specimen, make them unviable, or clog the surface or matrix due to wetting.

Moreover, many sensing formats, including ELISAs, bead-based assays, and other fluorescence- and thermophoresis-based techniques for biomolecular sensing, as outlined above, do not have the ability to directly extract absolute target protein concentrations from the recorded signal. Single-molecule detection can overcome this limitation, however, many sensing methods available to date are limited to bulk detection, and existing single-molecule sensing techniques often require the use of high-affinity probes, which are often hard to generate. For example, fluorescence correlation spectroscopy, requires the probe to bind at picomolar concentrations to enable quantitative binding and hence typically does not work for protein sensing when only a small proportion of probe is bound due to high background. A further challenge in traditional protein sensing formats is that many targets are usually required to be bi-epitopic, as detection is based on multiple binding probes. This increases design complexity because validated, non-cross-reactive affinity probe pairs are required. Furthermore, with many assays multi-step washing protocols for the removal of unbound probe from analyte are required, introducing lengthy workflows, on the order of tens of minutes to hours, which can cause analytes to dissociate, thereby limiting sensitivity.

Finally, and importantly, conventional techniques often do not inform on the nature of the analyte binding in terms of stoichiometry or immunochemistry/valency. Such multimodal information, however, is important, in particular for cases where it is relevant whether the species detected is monomeric or part of a complex (e.g., in pathologically relevant species such as amyloidogenic oligomers or fibrils) or where its expression levels need to be determined and the number of proteins on an exposed surface of the analyte may vary (e.g., of exosomal proteins). Taken together, sensing methods that avoid surface-capture of targets, have the ability to directly extract absolute target protein concentrations, and are based on a minimal number of steps for the removal of unbound probe in a rapid manner, yet increase the information content in terms of multimodal readouts are of increasing interest.

To address this challenge, we present here a single-molecule protein sensing approach, termed digital immunosensor assay (DigitISA), that operates in free solution and allows for second timescale separation of excess binding probe in a single, solution-phase step, allowing protein targets to be detected and their abundance to be quantified in solution without the need of washing steps. DigitISA’s design principle is based combining in-solution analyte capture in a surface-free manner and wash-free removal of unbound probe, thereby releasing fundamental constraints of conventional immunosensing approaches in terms of the thermodynamic and kinetics of the immunoprobe–analyte interaction. We achieve this objective by exploiting microfluidic free-flow electrophoretic separation, which allows protein-bound affinity reagents to be discriminated from non-protein bound ones based on a difference in their electrophoretic mobility (*i.e.*, the ratio of the net electrical charge of a molecule to its size). We combine the microchip electrophoretic separation step with in-situ single-molecule detection by laser-induced fluorescence confocal microscopy, which allows the number of protein-bound affinity reagent molecules present in the sample to be detected through single-molecule digital counting. Moreover, due to the digital nature of the detection process, the assay enables multimodal characterization of the binding interaction including immunochemistry/valency, binding stoichiometry, protein biomarker abundance, and provides further quantitative information of the probe–analyte interaction and composition.

In the following, we introduce the optofluidic DigitISA platform and detail its working principles and experimental implementation. We establish and validate the DigitISA assay by first probing a biomolecular biotin–streptavidin binding complex and demonstrate its applicability to biomedical analysis by quantifying IgE–aptamer binding. We show that DigitISA achieves in-solution single-molecule protein detection and quantification with picomolar sensitivity. We further use DigitISA to detect the presence of α-synuclein fibrils, a biomarker for Parkinson’s disease, using a low-affinity aptamer at high probe concentration. We then showcase label-free detection and analysis of α-synuclein oligomers, neurotoxic molecules and biomarkers in Parkinson’s disease, and provide information on oligomer immunochemistry by affording an estimate of the valency of aptamer–oligomer binding. We further demonstrate the ability of the DigitISA technique to sense and quantify proteins in complex, multi-component systems. We first probe the clinically relevant CD63 protein biomarker present on the surface of exosomes and simultaneously quantify the concentration of exosomes along with the expression levels of CD63, features that are hard if not impossible to obtain by other conventional means. Finally, we showcase the possibility to sense biomolecular condensates that are formed through liquid–liquid phase separation. Using an aptamer raised against the condensate forming protein fused in sarcoma (FUS), we demonstrate that the DigitISA platform can simultaneously determine absolute numbers of condensates in solution, determine concentrations of native protein in condensates and their mass densities, detect the stochiometric ratio of aptamers per condensate, determine the partitioning coefficient of aptamer into protein condensates, and provide information on the affinity of the monomer– aptamer interaction. Overall, our experiments demonstrate the versatility of the DigitISA platform in enabling multimodal in-solution quantification of protein biomarkers, including targets that would otherwise not be addressable by conventional immuno-sensing techniques.

## Results

### Working principle and assay design

The working principle and experimental implementation of the DigitISA platform is depicted in Figure 1a–c. Sample containing the target protein and a fluorescently labeled capture probe (e.g., aptamer) is injected into the microfluidic free-flow electrophoresis separation chip fabricated in poly(dimethylsiloxane) (PDMS) (see Methods: DigitISA platform and Fabrication of the electrophoretic devices). The sample stream, upon its entry into the separation region, is surrounded by a buffered carrier medium, so that the sample forms a narrow stream at the center of the chamber (Figure 1a,b). To discriminate the free probe from its analyte-bound form, an electric potential is applied perpendicular to the flow direction (Figure 1a) using co-flowing electrolyte solutions (Figure 1b) that act as liquid electrodes and ensure the application of stable electric fields, as described earlier.^19, 20^ Simultaneously, laser-induced confocal fluorescence microscopy is used to detect individual molecules in situ by scanning the confocal volume across the microfluidic chip (Figure 1a,c). Notably, scanning is performed in a stepwise manner at the mid-height of the channel at a distance of 4 mm downstream from the position where the sample first entered the electric field, as illustrated by the cyan line (Figure 1b). The number of molecules traversing the confocal volume at each of the scanned positions is estimated from the recorded photon-count time trace using a combined inter-photon time (IPT) and photon-count threshold burst-search algorithm (Figure 1a (see Methods: Data analysis). This approach has been shown to enable effective discrimination between photons that originate from single fluorescent molecules and those that correspond to a background,^21, 22^ thus allowing individual molecules to be counted directly, that is, in a digital manner. From the detected number of molecules at each position, an electropherogram is generated by plotting the obtained single-molecule counts as a function of chip position (Figure 1a). Thereby, the separation between protein-bound affinity probe and free probe can be visualized, and the concentration of the target complex quantified.

**Figure 1:**
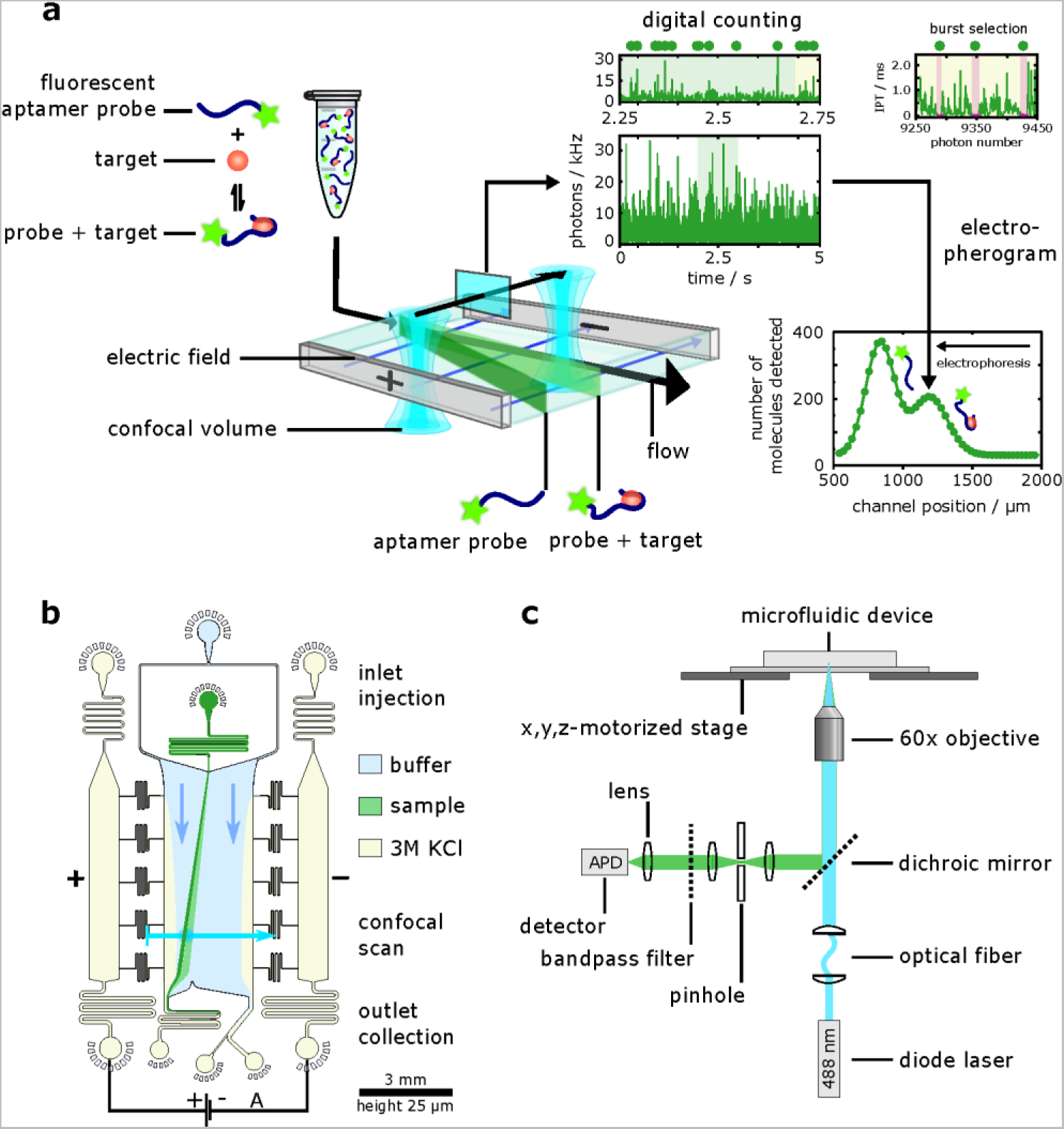
Working principle of DigitISA and its implementation. **(a)** Schematic illustration of the DigitISA platform and the experimental workflow. DigitISA integrates electrophoretic separation and single-molecule detection in a platform for single-step sensing of target proteins in solution using only a single affinity reagent. The sample including a mixture of the target protein and its fluorescently labeled affinity probe (e.g., aptamer) is injected into a micron-scale electrophoretic separation unit. The application of an electric field allows protein-bound probe molecules to be discriminated from those probe molecules that are not bound to the protein target, owing to a difference in their electrophoretic mobilities. Confocal scanning across the separation chamber is performed and the number of molecules traversing the confocal volume at each of the scanned positions is estimated ‘digitally’ from the recorded photon-count time trace using a combined inter-photon time (IPT) and photon-count threshold burst-search algorithm. From the obtained counts an electropherogram is created allowing for a discrimination between protein-bound affinity probe and free probe. **(b)** Design of the free-flow electrophoresis device. Sample is flown into the microfluidic chip by the central injection port where it is then surrounded by the carrier buffer solution. The electrophoresis chamber is connected to a co-flowing electrolyte solution (3 M KCl) via bridges, which allows for a narrow sheet of electrolyte to flow along the sides of the chamber. An electric field is applied from metal clips at the outlets of the electrolyte channels, which propagates along the electrolyte sheet and enables separation of molecules perpendicular to the flow direction. **(c)** Schematic of the confocal microscopy setup used for single-molecule detection. A diode laser is used to excite the sample through an objective, and a single-photon counting avalanche photodiode (APD) is used to register emitted photons from the sample. The confocal volume is moved across the cross-section of the chip in a stepwise manner with the aid of a motorized stage. This allows the flux of the protein-bound probe molecules to be estimated. Details of the setup are described in the Methods section (Data analysis).

### Sensing of a streptavidin–biotin affinity complex

To evaluate the possibility of using the combined free-flow electrophoresis and single-molecule counting DigitISA platform for biomolecular detection and quantification, we set out to probe the formation of a biotin–streptavidin complex. Specifically, we investigated the binding of a biotinylated and fluorophore-conjugated DNA sequence to monovalent streptavidin (see Methods: Protein preparation and Sample preparation for DigitISA experiments) (Figure 2a). This interaction mimics the binding of a protein molecule to its affinity reagent with the binding interaction being very well defined and of high affinity.^23^

**Figure 2:**
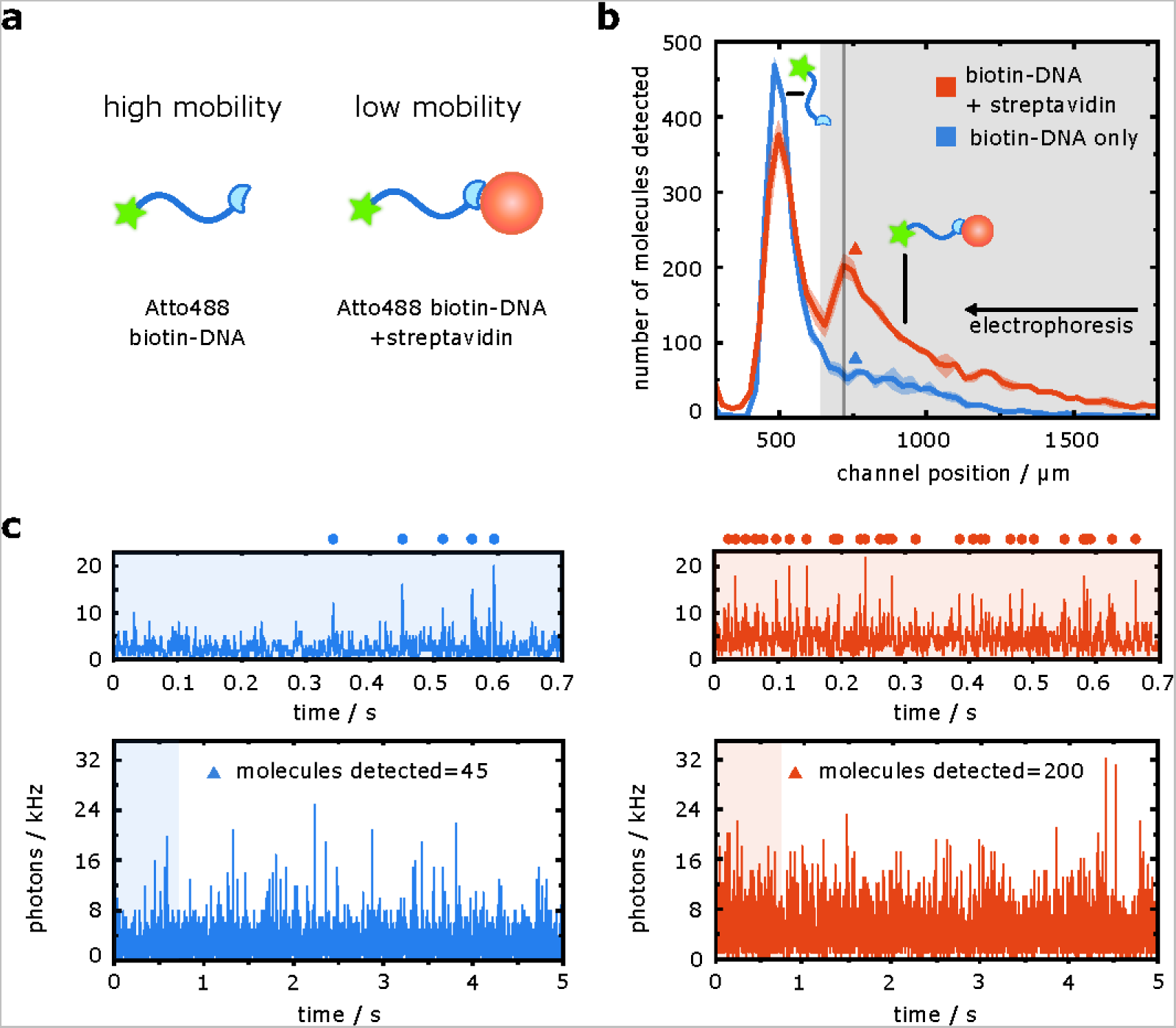
Sensing of a biotin–streptavidin complex using the DigitISA platform. **(a)** Monovalent streptavidin is added to a biotinylated and fluorophore-conjugated NA strand. Binding of the streptavidin species reduces the electrophoretic mobility of biotinylated DNA probe. **(b)** Electropherogram as obtained by scanning the confocal volume across the cross-section of the channel in a stepwise manner for the biotin–streptavidin sample mixture (red line; average of *N* = 3 repeats, the shaded bands correspond to the standard deviation) and for the control sample (blue line; average of *N* = 3 repeats, the shaded bands correspond to the standard deviation) at the mid-height of the channel, demonstrating the presence and separation of both streptavidin-bound and non-bound biotinylated DNA molecules in the sample. The region shaded in grey was used to extract the number of streptavidin–biotin complexes that passed the device in a given time, and ultimately, its concentration (see main text). **(c)** Exemplary photon-count time traces for the control sample (left panel, blue) and the sample mixture (right panel, red) at the position where the concentration of the complex molecules was the highest as indicated with colored triangles in panel b. The number of molecules was estimated using a burst-search algorithm as detailed in the Methods section (Data analysis). Time traces in the upper panels are zoom-in views of the blue or red shaded areas in the lower panels, with dots indicating detected single-molecule events. The bin time was 1 ms in all traces. Source data are provided as a Source Data file.

We first examined whether an applied electric field allows for a discrimination between the streptavidin-bound and unbound DNA probe according to a difference in electrophoretic mobility (Figure 2b). To this end, we incubated 25 pM of the monovalent streptavidin with 50 pM of the biotinylated probe DNA and injected the sample into the free-flow electrophoresis chip (see Methods: Experimental procedures) by applying an electric potential of 150 V, leading to electric fields of around 150 V cm^−1, 19, 24, 25^ which ensured that the biotinylated DNA probe deflects by a few hundred micrometers, allowing their discrimination from the streptavidin– biotin complex. Additionally, we injected a control sample including only the biotinylated DNA with no streptavidin into a different identically fabricated chip. 5-second-long step-scans along the cross-section of the microfluidic separation chamber were performed for both samples, and the number of molecules traversing the confocal volume at each of the scanned positions was estimated by performing a burst analysis on the photon time traces (Figure 2c, see Methods: Data analysis). Using the obtained single-molecule counts, electropherograms across the cross-section of the separation chamber were obtained for the two samples (Figure 2b) from *N* = 3 repeats. From these data, we observed that the binding of streptavidin decreased the electrophoretic mobility of the DNA-conjugated biotin molecules in comparison to the free DNA-conjugated biotin probe, with the free biotin eluting at a channel position of *x* = 500 µm and the streptavidin–biotin complex at *x* = 750 µm (Figure 2b, red line). Indeed, at the former position, only a minimal elution of fluorescent molecules occurred for the control sample (Figure 2b, blue line), indicating that this elution position corresponds to that of the biotin– streptavidin complex. Moreover, we also noted that the recorded fluorescence at the position where the unbound biotin–DNA molecules eluted was higher in the control sample than for the case when the streptavidin target was present. This observation further confirmed the integration of biotinylated DNA molecules into the complex.

Having confirmed the capability of the platform to detect the formation of the biotin– streptavidin complex, we next set out to explore its potential to estimate absolute concentrations. Indeed, while many protein-quantification assays, including conventional ELISAs, rely on a calibration curve between the observed signal and the concentration of the analyte molecules to extract absolute concentrations, the platform developed here counts the number of passing molecules in a digital manner (see Figure 2c for examples), and therefore has the potential to estimate target concentrations directly.

To explore this opportunity, we evaluated the number of molecules that eluted between channel positions of *x* = 700 µm and *x* = 2200 µm (Figure 2b, region shaded in grey). We recorded *n̄*_sample_ = 2391 ± 37 molecules (mean (*n̄*) ± standard deviation (*s*) of *N* = 3 repeats) for the biotin–streptavidin sample mixture (Figure 2b, red line) in contrast to only *n̄*_control_ = 789 ± 30 molecules for the control sample (Figure 2b, blue line) from *N* = 3 repeats. The non-zero count for the control sample is likely to originate from impurities in the biotin– DNA sample or from degraded forms of the DNA. The difference between the two counts, *n̄*_complex_ = *n̄*_sample_ – *n̄*_control_ = 1601 ± 48 molecules, can be attributed to streptavidin–biotin complexes. We note that this count corresponds to the molecular flux in the regions where the single-molecule time traces were recorded. The flux of the streptavidin–biotin complex molecules through the full device, *F*_total_, can be estimated from the following relationship:

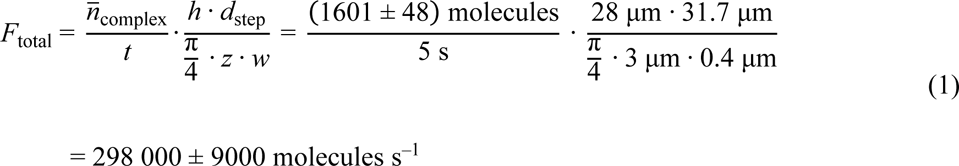

where *t* is the time period over which the time traces were recorded, *h* is the height to which the separation chamber was fabricated, *d*_step_ is the step size at which the single-molecule time traces were recorded, and *z* and *w* are the width and the height of the confocal detection volume, respectively, describing its cross-section. The latter two parameters were estimated from a fluorescent correlation spectroscopy (FCS) measurement (see Methods: Dimensions of the confocal volume from FCS) and were determined to be *z* = 3 µm and *w* = 0.4 µm for our setup.

As the sample was entering the device at a flow rate of *Q*_sample_ = 70 µL h^−1^, this molecular flux yielded an estimate of

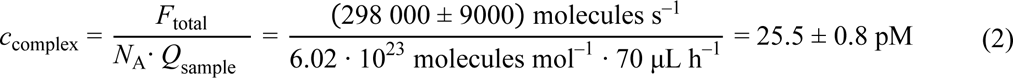

for the concentration of the streptavidin–biotin complex in the sample, with *N*_A_ being the Avogadro constant. The binding affinity between the monovalent streptavidin and biotin molecules has previously been estimated to be in the femtomolar range.^23^ As such, under the conditions used here (25 pM streptavidin and 50 pM biotin), we would indeed expect all of the monovalent streptavidin in the sample mixture to be incorporated into the complex. The recovery of the expected nominal complex concentration for this example thus validates our approach of direct digital counting and demonstrates that the platform can be used to obtain direct readouts of molecular concentrations.

### Immunosensing of the protein biomarker immunoglobulin E

Having established and validated the DigitISA platform for sensitive detection and quantification of biomolecular complexes, we set out to demonstrate the advantage of the fast (i.e., ∼2 s) assay timescale in the context of quantitative protein biomarker sensing. We investigated the protein immunoglobulin E (IgE), which is a key component of the human immune system. IgE shows a particular relevance in allergic responses, and elevated IgE concentrations are a defining characteristic of hyper-IgE syndrome and IgE myeloma.^26, 27^ We used an established IgE aptamer^28^ labeled with Atto488 fluorophore (see Methods: Sample preparation for DigitISA experiments) to detect the presence of IgE protein molecules (see Methods: Protein preparation) (Figure 3a). Aptamers are considered an attractive class of affinity reagents, whose production through in vitro evolution^29–31^ and chemical synthesis is fast and inexpensive, and they offer recognition capabilities that rival those of antibodies. Crucially, however, most conventional aptamers comprised of natural nucleotides (as opposed to the hydrophobically-modified bases employed in SOMAmer reagents)^14, 32^ are limited in binding strength, with typical values of *K*_d_ > 1 nM. Thus, quantitative sensing by aptamer probes in surface-based immunoassays is hindered by the relatively fast rate of probe–analyte dissociation, which has limited their widespread use, despite their many advantages. For example, with the IgE aptamer employed here (*K*_d_ ≈ 50 nM),^33, 34^ during an experiment time of 45 minutes as required for a ‘fast’ aptamer-based ELISA assay,^35^ approximately 66% of aptamer–analyte complexes present in the initial aptamer–analyte binding equilibrium would dissociate and not contribute to the sensing signal. Since DigitISA operates more than two orders of magnitude faster than ELISA-based approaches, negligible probe–analyte dissociation occurs on the assay timescale. Thus, the quantity of the aptamer–probe complex, probed by DigitISA, accurately reflects the equilibrium concentration of the complex present in the original probe–analyte mixture. Therefore, for an affinity probe of known *K*_d_, our method allows direct quantitative protein sensing in a single measurement, even at concentrations far below the *K*_d_ of the interactions.

**Figure 3:**
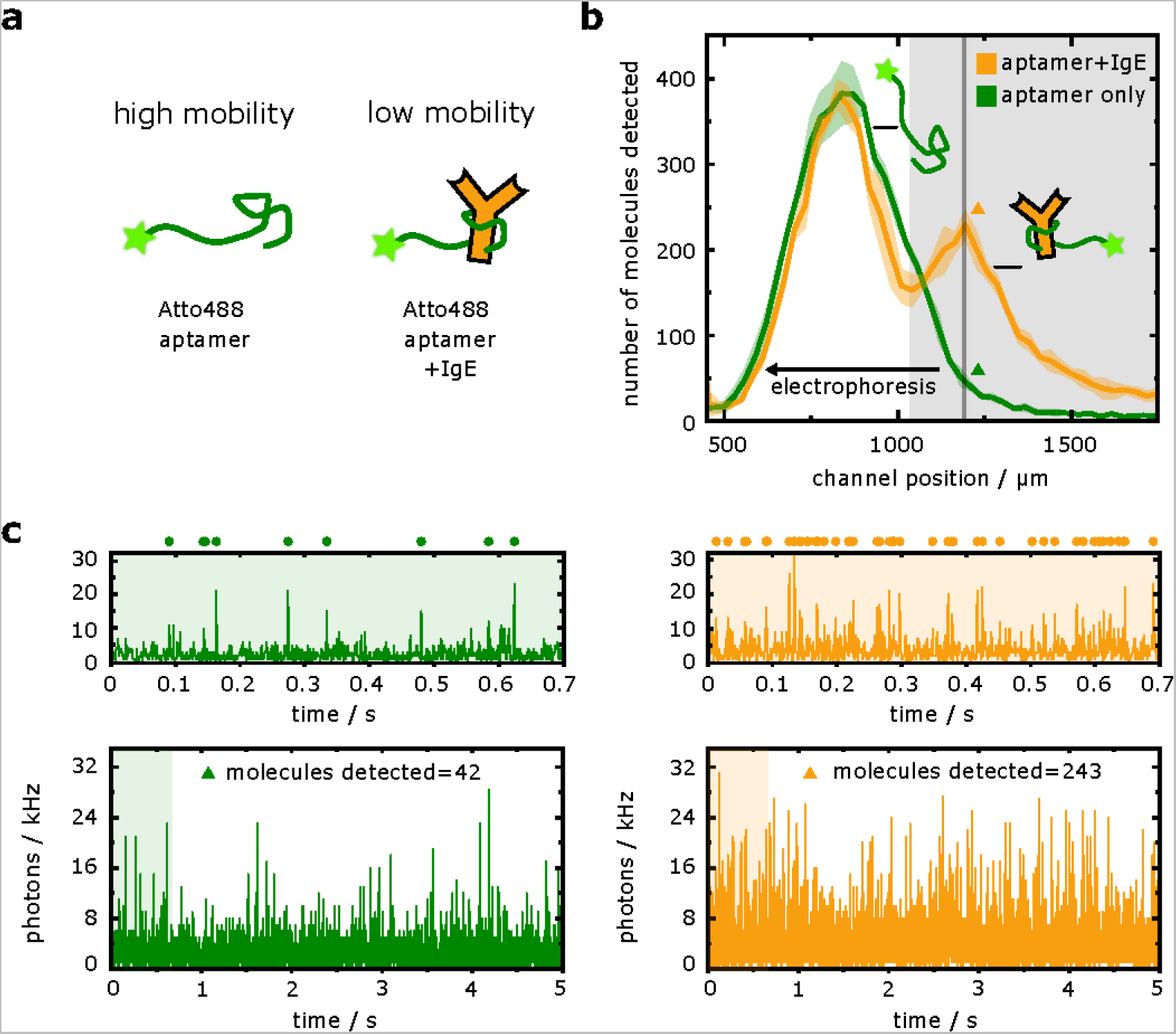
DigitISA immunosensing of IgE with an aptamer probe. **(a)** An IgE-aptamer probe is added to its target IgE. Binding of the antibody reduces the electrophoretic mobility of the probe, allowing for fast electrophoretic separation of the aptamer probe from the immuno-complex for subsequent confocal detection. **(b)** Electropherogram as obtained by scanning of the confocal volume across the cross-section of the channel in a stepwise manner for the IgE– aptamer sample (orange line; average of *N* = 3 repeats) and for the free aptamer probe (green line; average of *N* = 3 repeats). The shaded bands correspond to the standard deviation. The region shaded in grey was used to quantify the concentration of IgE by monitoring the flux of fluorescent molecules in the region shaded in grey (see main text). **(c)** Exemplary photon-count time traces for the control sample (left panel, green) and the sample mixture (right panel, orange) at the position where the concentration of the complex molecules was the highest as indicated with colored triangles in panel b. The number of molecules was estimated using a burst-search algorithm as detailed in the Methods section (Data analysis). Time traces in the upper panels are zoom-in views of the green or orange shaded areas in the lower panels, with dots indicating detected single-molecule events. The bin time was 1 ms in all traces. Source data are provided as a Source Data file.

To demonstrate this principle, IgE (40 nM) was premixed with aptamer probe (50 pM) and injected into the microfluidic separation device, and electropherograms were acquired (Figure 3b, orange line) in a similar manner to the streptavidin–biotin system described above (from *N* = 3 repeats) (see Methods: Experimental procedures). The concentrations were chosen so that only a small proportion of protein would be bound by the probe, to test whether our approach would allow effective back-calculation of the true protein concentration. As expected, both the free probe and the probe–protein complex were observed in the electropherograms with the complex eluting at a higher channel coordinate (smaller deflected distance) due to its reduced mobility (Figure 3b, orange line). Control experiments with no IgE showed only the free-probe peak with a minimal amount of fluorescence detected at the elution position of the complex (Figure 3b, green line). As before, we used the photon-count time traces (from *N* = 3 repeats) (see Figure 3c for examples) to estimate the flux of fluorescent molecules in the regions where the complex eluted (Figure 3b, region shaded in grey). From these data, we concluded that *n̄*_sample_ = 2847 ± 62 molecules eluted over a time period of 5 s for the sample mixture and *n̄*_control_ = 856 ± 29 molecules for the control sample that included only the probe molecules without the target. Using a conversion strategy similar to what was described above for the biotin–streptavidin system, this molecular flux value yielded an estimate of *c*_complex_ = 21.7 ± 0.7 pM for the concentration of the aptamer–IgE complex.

Using a simple 1:1 binding model for the aptamer–IgE interaction and our knowledge of the concentrations of the IgE bound (*c*_complex_ = 21.7 pM) and free (*c*_free_ = 50 pM– 21.7 pM = 28.3 pM) forms of the aptamer probe, we calculated the concentration of IgE present in the sample to be *c*_IgE_ = 38.4 ± 1.6 nM according to *c*_IgE_= *c*_complex_·*K*_d_⁄*c*_free_. This value showed excellent agreement with the nominal starting concentration of IgE, thereby validating our approach of direct digital counting in this concentration range, with the small deviation likely originating from an uncertainty in the reported value of the binding constant *K*_d_.^33, 34^ This result demonstrates the efficacy of our approach for quantitative protein sensing even for relatively weak-binding probes, as enabled by the rapid fractionation of protein bound and unbound probe by the microchip free-flow electrophoresis platform, and provides a route to performing measurements in a regime where the availability of the affinity reagent is limited. Notably, our approach exploits the fact that detection relies only on a single, monovalent interaction between the probe and the analyte, which allows facile back-calculation of the target concentration from the underlying binary equilibrium, as demonstrated above. Conversely, even for a well-characterized ELISA experiment, such calculation is challenging to realize given the multiple analyte–antibody equilibria that are present in the sandwich-type formats employed in ELISAs.

### Sensing of α-synuclein amyloid fibrils

Having shown the advantages associated with the fast assay timescale, we next demonstrated the ability of the free-solution DigitISA assay to directly sense the probe–analyte binding equilibrium by the use of high probe concentration. To do so, we investigated the binding between α-synuclein fibrils and an aptamer (see Methods: Protein preparation and Sample preparation for DigitISA experiments), which has been shown previously to weakly bind fibrillar forms of the α-synuclein protein with an approximate *K*_d_ of 500–1000 nM.^36^ Fibrillar α-synuclein is a molecular hallmark of Parkinson’s disease and other synucleinopathies; sensing of α-synuclein aggregates is thus proposed as a means for the early detection for these conditions.^37^ As illustrated in Figure 4a, binding of the aptamer to the α-synuclein fibrils is expected to suppress the electrophoretic mobility of the fibrils, thus enabling efficient separation of aptamer-bound fibrils from the unbound probe that is provided in excess.

**Figure 4:**
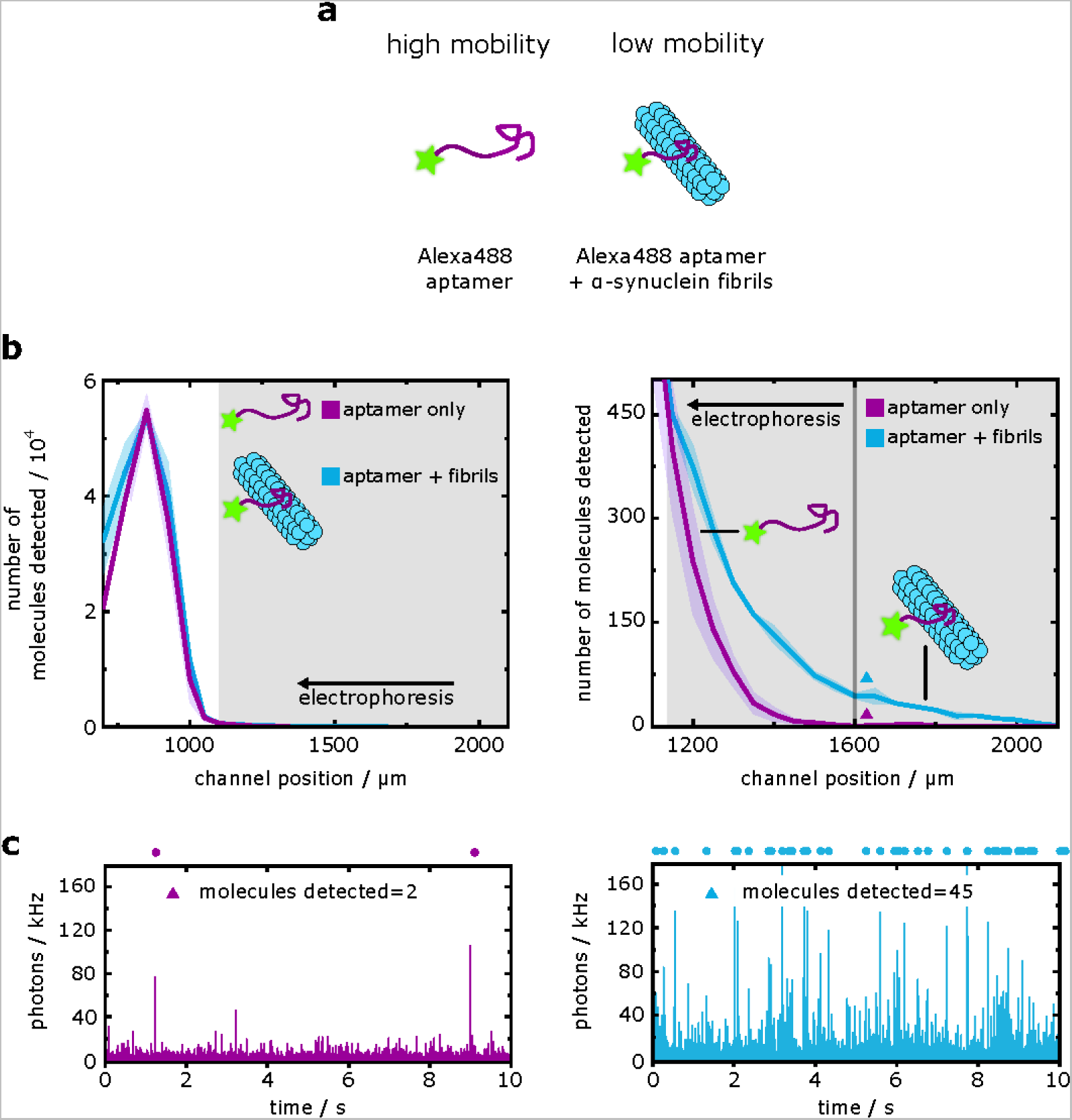
Sensing of α-synuclein fibrils at high aptamer probe concentrations. **(a)** Binding of an α-synuclein aptamer to fibrils reduces the electrophoretic mobility of the aptamer probe, allowing for the discrimination between fibril-bound and unbound aptamer species. **(b)** Electropherogram (left panel) as obtained by scanning of the confocal volume across the cross-section of the channel in a stepwise manner for the α-synuclein fibrils–aptamer sample (blue line; average of *N* = 3 repeats) and for the free aptamer probe (purple line; average of *N* = 3 repeats). The shaded bands correspond to the standard deviation. The right panel shows a zoom-in region of the electropherogram. The shaded region in grey where the concentration of the complex exceeded that of the free probe (1150 µm < *x* < 2000 µm) was used to estimate the concentration of the fibrils (see main text). Note, the photon arrival frequency in the region where *x* < 1100 µm (i.e., where the probe elutes) was too high to count molecules one-by-one, hence the detected number of molecules in this region should be viewed as an approximation. **(c)** Exemplary photon-count time traces for the control sample (left panel, purple) and the sample mixture (right panel, blue) at the position indicated with colored triangles in panel b (right). The number of molecules was estimated using a burst-search algorithm as detailed in the Methods section (Data analysis). Time traces in the upper panels are zoom-in views of the purple or blue shaded areas in the lower panels, with dots indicating detected single-molecule events. The bin time was 1 ms in all traces. Source data are provided as a Source Data file.

To demonstrate this capability, we incubated a sample of the fibrils with 10 nM of aptamer probe and acquired electropherograms (from *N* = 3 repeats) by applying an electric potential of 80 V (Figure 4b, blue line) (see Methods: Experimental procedures). By comparison of the averaged electropherograms for the aptamer–fibril sample relative to the aptamer-only controls (Figure 4b, purple line), fibril–aptamer complexes could be identified at lower electrophoretic mobilities relative to the unbound aptamer peak. We estimated there to be a total of *n̄*_sample_ = 2066 ± 47 molecules for the aptamer–fibril sample mixture and *n̄*_control_ = 926 ± 132 molecules of the unbound aptamer eluting in the shaded grey region highlighted in Figure 4b (1150 µm < *x* < 2100 µm) over a 10-second timescale (see exemplary time traces in Figure 4c). Using a similar analysis as before, these molecular counts yielded the concentration of protein-bound probe to be 18.7 ± 2.3 pM for the α-synuclein fibrils. The concentration of protein-bound probe corresponds to a total fibril binding site concentration of 1.0–1.9 nM for a *K*_d_ range of 500–1000 nM.

We note that for a theoretical ELISA experiment with the same analyte concentration and a probe concentration of 1 nM, approximately only 1.9–3.8 pM of fibril would be captured on the surface. Subsequent, rapid dissociation of the probe–aptamer complex due to the weak probe binding strength over the assay timescale would reduce the concentration of bound target well below the assay detection limit. Together, these factors illustrate how the DigitISA platform can be used for the detection of weak biomolecular binding interactions—a characteristic that is challenging to achieve with conventional multi-step approaches operating over longer time scales.

### Label-free detection and analysis of α-synuclein oligomers

In addition to sensing low abundance αS fibrils, our DigitISA technique can also be applied to the sensing of αS oligomers. These assemblies are considered to be the main pathogenic species causing neurodegeneration in Parkinson’s disease.^38, 39^ A challenge still remains to find methods to detect these low abundance species and especially to be able to sense wild-type oligomers without fluorophores attached.

To demonstrate the sensing of αS oligomers by DigitISA, we set out to probe lowly populated and transient oligomeric species formed when the αS protein aggregates (see Methods: Protein preparation and Sample preparation for DigitISA experiments). We employed an oligomer-selective, fluorescently-labelled aptamer probe, which has good affinity for αS oligomers (*K*_d_ = 150–200 nM) but does not bind αS monomer.^36^ Aptamer–protein complexes have been shown to have reduced electrophoretic mobility relative to unbound aptamer, enabling electrophoretic separation of the free and protein-bound probe.^24, 40^ We incubated an aliquot taken from a stirring-induced aggregation of wild-type αS and added an excess of aptamer probe (100 nM) to enable efficient oligomer capture (Figure 5a) (see Methods: Experimental procedures). Oligomer-bound aptamers (Figure 5b, blue line) were indeed separable from the excess aptamer probe (orange line), enabling single-molecule counting of oligomer–aptamer complexes as they transited the confocal volume in DigitISA experiments. From *N* = 3 repeats, we estimated there to be a total of *n̄*_sample_ = 1532 ± 439 molecules for the aptamer–oligomer sample mixture and *n̄*_control_ = 665 ± 247 molecules of the unbound aptamer eluting in the shaded grey region highlighted in Figure 5b (1000 µm < *x* < 2100 µm) over a 30-second timescale (see exemplary time traces in Figure 5d). Converting from molecule counts to concentration (Eq. 1 and 2), a concentration of 32.7 ± 7.2 pM could be estimated for the sample of wild-type oligomers. The ability of DigitISA to provide absolute concentrations of αS oligomers in a label-free manner is significant, since alternative measures of oligomer concentration given by radiolabelling^41^ or UV-VIS absorption^42^ only provide mass concentration, and cannot elucidate the number of individual oligomer species as is enabled by our digital-counting platform.

**Figure 5:**
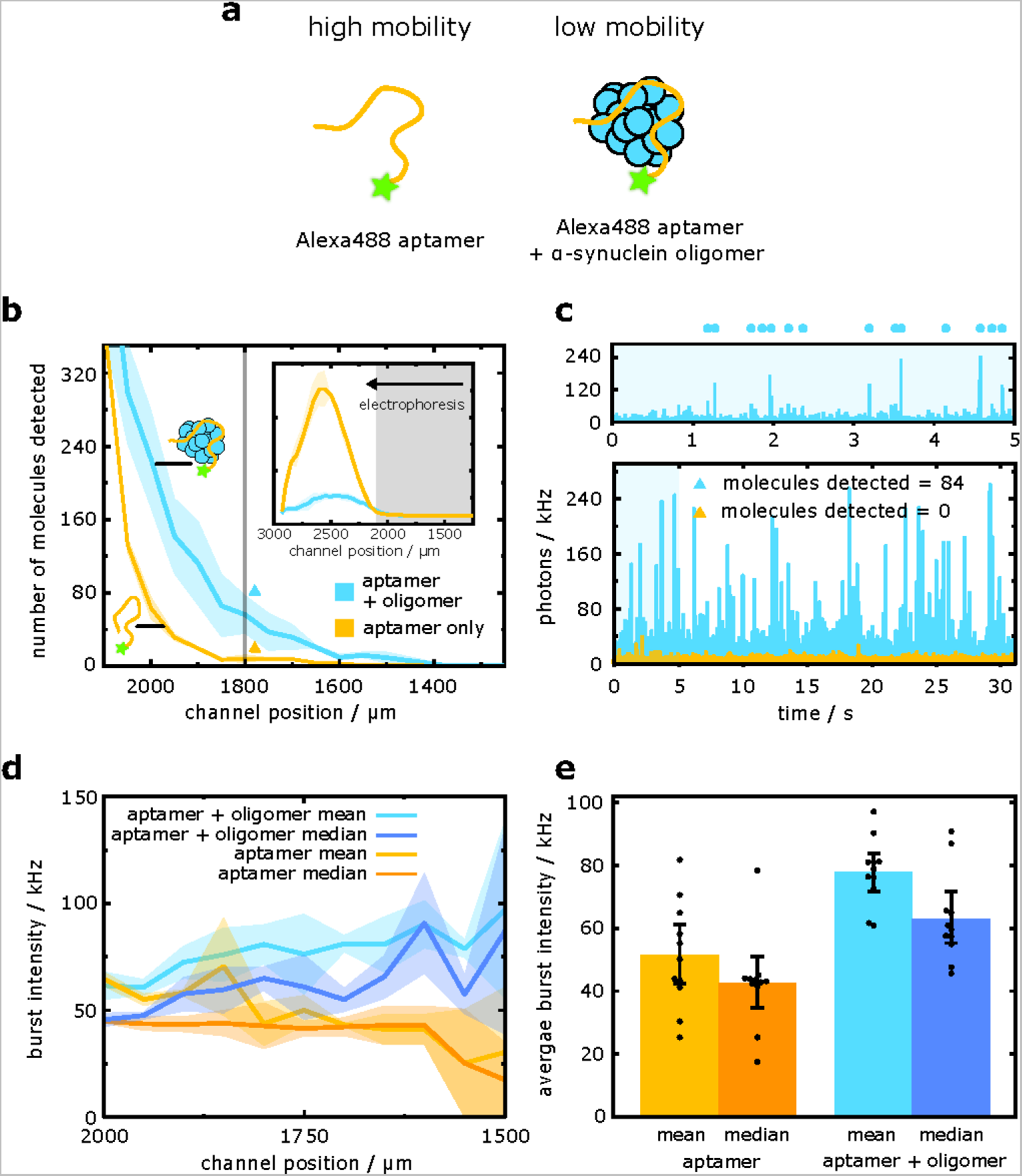
Sensing and immunochemical analysis of α-synuclein oligomers. **(a)** The binding of an aptamer to wild-type α-synuclein oligomers reduces the electrophoretic mobility of the probe, allowing for the discrimination between oligomer-bound and unbound aptamer species. **(b)** Electropherogram as obtained by scanning of the confocal volume across the cross-section of the channel in a stepwise manner for the α-synuclein oligomer–aptamer sample (blue line; average of *N* = 3 repeats) and for the free aptamer probe (orange line; average of *N* = 3 repeats). The shaded bands correspond to the standard deviation. The main panel depicts the region where single-molecule counting of the probe or oligomer–probe complex is possible. This region was used to estimate the concentration of oligomers in the sample. The inset shows the full electropherogram and the average photon intensity at each step location, with the region shaded in grey depicting the region shown in the main panel. **(c)** Exemplary photon-count time traces for the control sample (orange) and the sample mixture (blue) at the position indicated with colored triangles in panel b. The number of molecules was estimated using a burst-search algorithm as detailed in the Methods section (Data analysis). Time trace in the upper panels is a zoom-in view of the blue shaded area in the lower panel, with dots indicating detected single-molecule events. The bin time was 1 ms in all traces. **(d)** Burst intensities of the aptamer– oligomer (blue) and free aptamer (orange) detection events across the single-molecule detection region of the channel. Both the mean and median intensities are reported. Each line is the average of *N* = 3 repeats, the shaded bands correspond to the standard deviation. **(e)** Average burst intensities of the free aptamer (orange) and aptamer–oligomer sample (blue) across the single-molecule detection region of the channel (see panel d). Both the mean and median intensities are reported. Error bars denote the standard deviation of *N* = 3 repeats of all data points across the single-molecule detection region of the channel (see panel d). Data points are shown. Source data are provided as a Source Data file.

In addition to quantifying the concentration of wild-type oligomers, further analysis can also yield information on oligomer immunochemistry, by affording an estimate of the valency of aptamer–oligomer binding. We noted that bursts of fluorescence corresponding to the passage of oligomer–aptamer complexes through the confocal spot were more intense than that of unbound aptamer (Figure 5d,e). This observation is suggestive of multivalent oligomer–aptamer binding. Based on the ratio of the averaged median burst intensities of *I*complex = 59.5 ± 13.6 and *I*_free aptamer_ = 43.0 ± 14.2 photons per burst, we estimate an approximate average of 1.38 aptamers bound to each oligomer. Of note, we used the median intensities in estimating the intensity ratio, as the median is resilient to bias from burst distribution shape. With the burst ratio information in hand, by considering the aptamer–oligomer affinity (*K*_d_ = 150–200 nM) and concentrations of oligomer-bound (*c*_complex_ = 32.7 pM) and free aptamer (*c*_free_ = 100,000 pM – (1.38 · 32.7 pM)= 99,954 pM), we approximate, using a simple model of aptamer–oligomer association, that the oligomer binding site concentration amounts to *c*_bindingsite_ = 82–98 pM, according to *c*_binding site_ = (*c*_complex_⁄*c*_free_)(*K*_d_+*c*_free_). Dividing this value by *c*_complex_ yields an apparent number of binding sites and shows that on average each oligomer possesses 2.5–3.0 aptamer binding sites per oligomer. Evidence for multivalent binding implies that the surfaces of αS oligomers may have multiple repeat units in oligomer quaternary structure and binding epitopes which generates the number of aptamer binding sites calculated here. These findings are particularly significant in the context of therapeutic efforts to ameliorate the effect of misfolding diseases, as they demonstrate that oligomers possess multiple binding sides that could potentially be targeted by immunotherapies developed against neurodegenerative proteinopathies.^43^

More generally, our findings demonstrate the capability of our platform to enable both highly sensitive number–concentration quantitation of wild-type oligomers and characterization of oligomer immunochemistry and stoichiometry in a single measurement. Notably, these observations are only achievable by the single-complex resolution inherent to our approach, and would be impossible using conventional immunochemical techniques such as ELISA, which would produce a population-averaged readout of aptamer binding without elucidating the true concentration of oligomers present or the number of aptamers bound per oligomer.

### Marker protein detection and quantification of exosomes

Next, to demonstrate the ability of the DigitISA technique to sense and quantify protein biomarkers in complex, multi-component systems, we set out to probe a biomarker present on the surface of exosomes. Exosomes are nanometer-sized extracellular vesicles of endosomal origin that carry a large number of proteins, RNAs and other biomolecules.^44^ They are ubiquitously found in body fluids and shed by cells under both physiological and pathophysiological conditions.^45, 46^ Exosomes are therefore considered important and clinically relevant diagnostic markers for a wide range of diseases, in particular for early cancer diagnosis and prognostics.^47–49^

To demonstrate exosome biomarker sensing by DigitISA, we set out to probe the exosomal surface marker protein CD63 (Figure 6a). CD63 is a member of the tetraspanin membrane protein family and used as a marker for predicting and monitoring the prognosis of cancers and other diseases.^50^ We used an aptamer developed against CD63 that has previously been employed in immuno-histochemistry and other biosensing applications (see Methods: Sample preparation for DigitISA experiments) (Figure 6a).^51^ We isolated exosomes from the breast cancer cell line MDA-MB-231 (see Methods: Exosome preparation), a widely used cell line in research on breast cancer metastasis and in drug development.^52^ We characterized exosomes with TEM imaging and used immunoblotting to detect the presence of CD63 (Figure 6b). The exosome isolates were mixed with fluorescently labeled aptamer, at three different concentrations (5 nM, 50 nM, and 100 nM), and injected into the microfluidic separation device (see Methods: Experimental procedures). Additionally, we performed a control experiment with aptamer only at 50 nM. All experiments were performed *N* ≥ 3 times. Exemplary electropherograms for the experiment performed at 50 nM aptamer concentration and the control experiment are shown in Figure 6c. Free aptamer and aptamer-bound exosomes could readily be separated based on their electrophoretic mobility, with the exosomes having lower mobility both due to their larger size and lower average charge relative to free aptamer. Free aptamer, in both the aptamer-only and aptamer–exosome sample, eluted at channel positions typically >2400 µm. Single-molecule analysis of the region at lower channel coordinates (i.e., lower mobilities) revealed an additional peak at channel coordinates around 2300 µm, which was absent in the control sample. Inspection of the time traces at these lower mobility positions (Figure 6d) showed fluorescence burst events of high intensity, stemming from aptamer-bound exosomes passing the detection volume, while none of such bursts were detected in the control sample.

**Figure 6:**
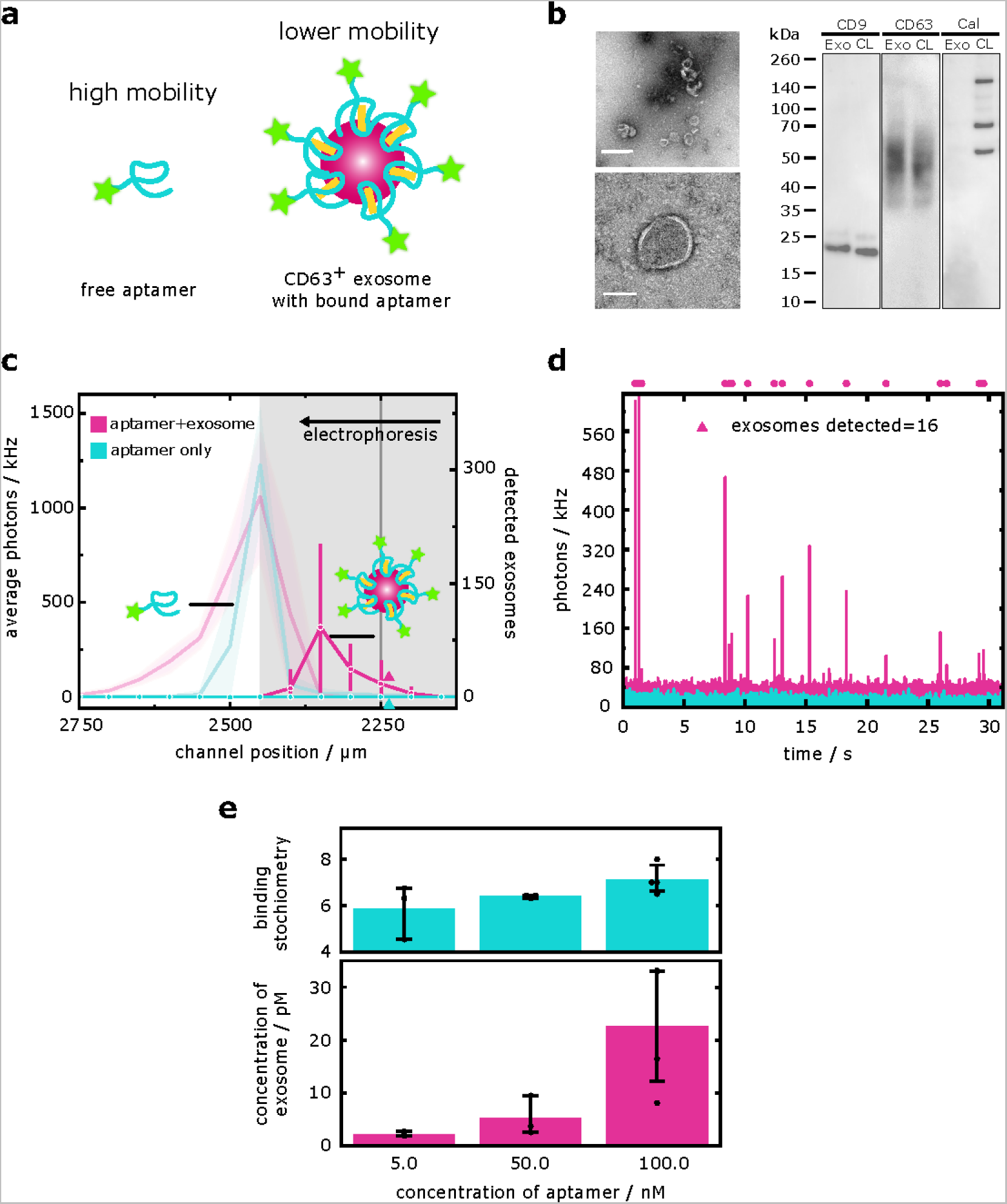
DigitISA immunosensing of an exosome biomarker. **(a)** Shown is a CD63-specific aptamer bound to CD63 proteins on the surface of the exosomes. The exosome-bound aptamers have a lower electrophoretic mobility than the free aptamers, allowing for the discrimination between exosome-bound and unbound species. **(b)** TEM micrographs (left panels) and Western blots (right panels) of exosomes. Exosomes display their characteristic cup-shaped morphology and can be identified by their size (100–200 nm in diameter) and the presence of a lipid bilayer membrane. TEM experiments were repeated three times. Scale bars: 500 nm (top left), 100 nm (bottom left). Western blot (right panels) comparing exosome samples (Exo) and cell lysate sample (CL). CD9 and CD63, both exosome marker proteins, are highly enriched in the exosome sample, while calreticulin (Cal) is excluded from the exosomes, indicating their purity. Western blot experiments were repeated three times. **(c)** Electropherograms as obtained by scanning of the confocal volume across the cross-section of the channel in a stepwise manner for the 50 nM exosome–aptamer sample (magenta line; average of *N =* 3 repeats) and for the free aptamer sample (cyan line; average of *N* = 3 repeats). The shaded bands correspond to the standard deviation. The light lines with error bands represent the average fluorescence intensity. The solid markers with error bars indicate the number of single-molecule detection events at each coordinate with the error bar representing the standard deviation. The region shaded in grey was used to estimate the concentration of exosomes in the sample. **(d)** Exemplary photon-count time traces for the control aptamer-only sample (cyan) and the exosome mixture (magenta) at the position indicated with colored triangles in panel c. Individual exosomes were detected and quantified using a burst-search algorithm as detailed in the Methods section (Data analysis). Dots above the time traces indicate individual exosome detection events. The bin time was 1 ms in all traces. **(e)** Mean concentrations of bound exosomes and averaged median binding stoichiometries at three different concentrations of aptamer. Error bars denote the standard deviation of *N* ≥ 3 repeats. Data points are shown. Source data are provided as a Source Data file.

Using a burst search algorithm, we extracted the number of exosomes detected in the regions corresponding to where aptamer-bound exosomes eluted (2000 µm < *x* < 2400 µm). This allowed us to calculate the concentration of exosomes that have the CD63 marker expressed on their surface. The number of detected CD63-positive exosomes (from *N* ≥ 3 repeats) increased with increasing aptamer concentration from *n̄*_exosomes, 5 nM_ = 88 ± 18 to *n̄*_exosomes, 50 nM_ = 210 ± 151 and *n̄*_exosomes, 100 nM_ = 915 ± 504 (Figure 6e). Converting from molecule counts to concentrations (Eqs. 1 and 2), the concentrations of detected CD63-positive exosomes were *c*_exosomes, 5 nM_ = 2.2 ± 4.1 pM, *c*_exosomes, 50 nM_ = 5.2 ± 3.7 pM, and *c*_exosomes, 100 nM_ = 22.7 ± 12.5 pM. This concentration series, with increasing amount of aptamer probe, shows the ability for DigitISA to assess binding curves at low analyte concentrations on the order of tens of pM, with high specificity, even in a heterogeneous extracellular mixture. This ability for DigitISA to be used as an affinity assay between low affinity probes and extremely dilute analytes may be helpful in cases such as the detection of exosomes in serum samples from cancer patients.^53^ In addition to digital counting of exosomes, the DigitISA technique allows us to extract the number of CD63 molecules present on the exosome surface (i.e., expression level). This is done by determining the individual burst intensity of each detected event and normalizing the burst intensity by the intensity of a single fluorescent aptamer. Averaged median burst intensities (from *N* ≥ 3 repeats) for the three exosome samples were *I*_exosome, 5 nM_ = 93.8, *I*_complex, 50 nM_ = 102.3, *I*_complex, 100 nM_ = 114 photons per burst; the averaged median burst intensity of the free aptamer (from *N* = 3 repeats) was *I*_free aptamer_ = 16.0 photons per burst. This yielded an averaged median number of aptamers bound per exosome in the range of 5.8–7.1 (Figure 6e), showing that the number of aptamers per exosome has little variation across the binding series. This value is in excellent agreement with previously reported binding stoichiometry values determined by flow cytometry with a CD63 specific antibody.^54^

In summary, our data on exosome biomarker detection shows that the DigitISA platform is well suited for the analysis of biomarkers in complex samples. We simultaneously quantify the concentration of exosomes along with the expression levels of CD63, features that are hard if not impossible to obtain by other conventional means. Considering the availability of other aptamers and alternative strategies for protein and RNA detection, this study opens a wide possibility of using DigitISA for the characterization of clinically relevant exosome samples and other complex multi-component systems.

### Sensing of biomolecular condensates

To further display the capabilities of the DigitISA platform, we explored the possibility to sense biomolecular condensates that are formed through liquid–liquid phase separation. Demixing of proteins and other biomolecules into liquid-like condensates has in recent years emerged as an underlying principle of subcellular organization^55^ and has become of particular interest because phase separation processes are implicated in diseases such as cancer^56^ and neurodegenerative disorders.^57, 58^ There is currently an unmet need in developing quantitative sensing assays for the detection of biomolecular condensates, as these assemblies are becoming important targets in drug discovery and diagnostics;^59^ yet, due to their liquid-like nature, they evade scrutiny by classical protein sensing methods. Phase-separated systems are inherently heterogenous because they demix into a protein-depleted diluted phase, consisting of monomeric protein, and a protein-enriched condensed phase consisting of micron-sized condensates. This makes it challenging to sense each phase state simultaneously, not least because of the largely different size and abundance of condensates and monomeric protein. In fact, a condensate system would not be addressable by conventional binding assay techniques (e.g., ELISAs), simply because it would be impossible to distinguish two different species by affinity probes as they would both contribute to the same signal. By contrast, because of its single-molecule sensitivity and combined physical separation capabilities, DigitISA is well-suited to tackle this challenge.

To demonstrate condensate detection by DigitISA, we set out to probe the phase behavior of the protein fused in sarcoma (FUS) (see Methods: Protein preparation). FUS is a DNA/RNA-binding protein that is involved in transcription regulation, RNA transport, and DNA repair,^60, 61^ and has been in the focus of many studies in recent years, due to its ability to phase separate, either on its own^62^ or in complex mixtures with nucleic acids,^63^ and its implications in neurodegenerative diseases and cancer.^64, 65^ To utilize the DigitISA technique for FUS detection, we used a fluorescently labelled aptamer (see Methods: Sample preparation for DigitISA experiments), that is known to have a mid-range affinity for the FUS protein in the hundreds of nanomolar regime.^66^ As illustrated in Figure 7a, binding of the aptamer to FUS is expected to suppress the electrophoretic mobility of the monomeric protein and of condensates. We phase separated FUS at a concentration of 2 µM and added 700 nM of fluorescently tagged aptamer, which resulted in the formation of condensates (Figure 7b). We injected the mixture into the microfluidic separation device (see Methods: Experimental procedures) and acquired electropherograms at 150 V by recording 5-second-long time traces across the device (from *N* = 3 repeats). The electropherograms exhibited two peaks (Figure 7c), corresponding to free probe aptamer at lower channel coordinates, and FUS–aptamer eluting at higher channel positions (i.e., smaller deflected distance) due to its reduced mobility. Control experiments (from *N* = 3 repeats) with no FUS showed only the free-probe peak, which was well-separated from the FUS–aptamer peak (Figure 7c). Inspection of the time traces (Figure 7d) revealed that the signal of the FUS–aptamer peak consists of a bulk signal, representing monomeric FUS, in addition to burst signals with high intensity, representing FUS condensates passing the confocal volume. The presence of an excess of FUS monomer over condensates is due to a reentrant transition of the condensate phase, due to which FUS can partially dissolve back from its condensate state to homogenous monomer in the presence of nucleic acids.^63, 67^

**Figure 7:**
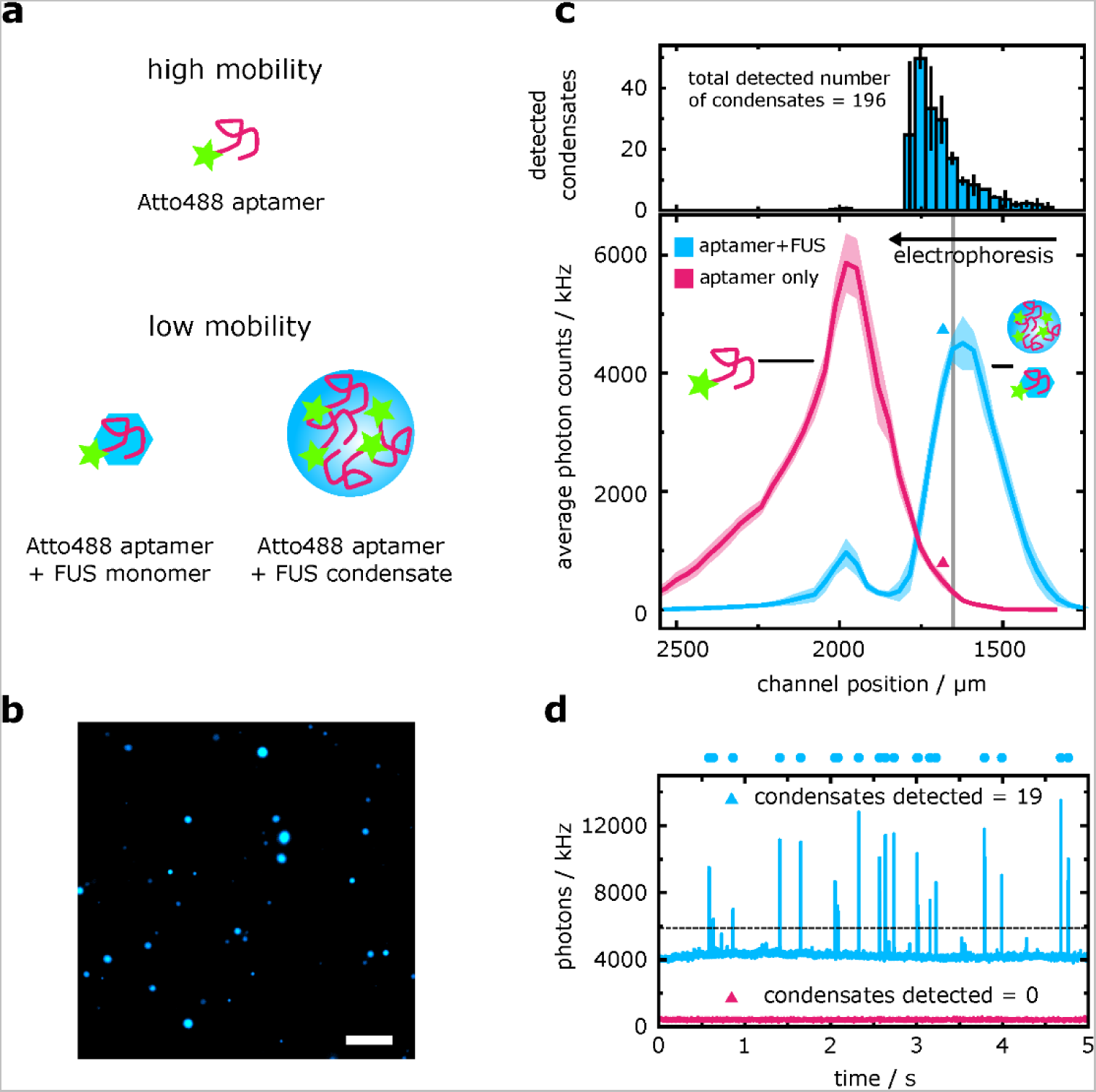
DigitISA sensing of protein condensates. **(a)** A low affinity aptamer for FUS protein can be found in three different states: unbound, with high electrophoretic mobility, bound to monomer protein, or partitioned into FUS condensates, the latter two exhibiting lower electrophoretic mobilities. **(b)** Fluorescently labelled GGUG aptamer is added to unlabeled FUS, resulting in the formation of biomolecular condensates. Image was taken on a widefield fluorescence microscope. Experiment was repeated three times. Scale bar is 10 µm. **(c)** Electropherogram as obtained by scanning of the confocal volume across the cross-section of the channel in a stepwise manner for the FUS–aptamer sample (blue line; average of *N* = 3 repeats, the shaded bands correspond to the standard deviation) and for the free aptamer probe (magenta line). Note, the ordinate shows the average fluorescence intensity, which was used to calculate the ratio between free and FUS monomer-bound aptamer (see main text). In the top panel, a histogram of the number of condensates detected at each channel position is shown, which was used to calculate condensate number and concentration. Error bars denote the standard deviation of *N* = 3 repeats. **(d)** Exemplary photon-count time traces for the control sample (magenta) and the condensate mixture (blue) at the position indicated with colored triangles in panel c (bottom). The number of condensates was quantified by counting bursts above a relative intensity threshold (shown by the dashed line) unique to each scan position, a process further detailed in the Methods section (Data analysis). Dots above the time traces indicate detected single-condensate events. The bin time was 1 ms in all traces. Source data are provided as a Source Data file.

Using these data, we set out to extract affinities of the aptamer for both the FUS monomer and FUS condensates. First, we calculated the fractions of monomer-bound and free aptamer by fitting a Gaussian curve to each of the bound and unbound peaks in Figure 7c. From this we determined the equilibrium binding of aptamer to FUS as *K*_d_ *=* 280 ± 20 nM, consistent with previous results.^66^ We further quantified the number density and volume fraction of condensates. By counting the number of bursts in the entire channel range where sample was present (1000 µm < *x* < 2600 µm) (Figure 7d), we concluded that a total of *n*_condensates_ = 196 ± 38 condensates eluded in the FUS–aptamer peak (Figure 7c, top panel). Applying the above analysis routine (Eqs. 1,2), this corresponds to a condensate particle concentration of *c*_condensates_ = 11.0 ± 2.2 pM, or more significantly, a volume fraction of *ϕ*_condensates_ = 0.5–2.5%, assuming an average radius of the condensates of *r*_condensates_ = 825 ± 454 nm (Figure 7b, taken from 400 detected condensates in widefield fluorescence imaging). Note that the condensates are highly polydisperse in size, which leads to a large standard deviation of the radius and subsequent variation in volume fraction.

To further analyze the aptamer–condensate interaction, we determined the partitioning ratio of the aptamer into the condensate. To this end, we compared the average fluorescence intensity of detected condensates (*I*_condensates_ = 7307 ± 2793 photons) with the average intensity of the free aptamer (*I*aptamer = 22.7± 0.9 photons), and obtained an apparent stoichiometry of 322 ± 123 aptamers per condensate. Multiplied by the determined concentration of condensates (*c*_condensates_ = 11 ± 2.2 pM), we find that the concentration of aptamer partitioned into condensates amounts to *c*_aptamer, FUS condensate_ = 3.5 ± 1.5 nM. In comparison, the amounts of aptamer associated with the monomer protein and free in solution are *c*_aptamer, FUS monomer_ = 586.5 ± 41.8 nM and *c*_free, aptamer_ = 110.0 ± 7.9 nM, respectively, as determined by the integral of the respective signal peaks in the electropherograms (Fig 7C). From this we find that 3.2% of the free RNA partitions into the condensates; if we additionally consider the RNA that is bound to FUS monomerically, then we determine 0.5% of the total aptamer concentration is present in condensates.

We further determined the protein concentration in the condensate phase. Assuming that all protein in the non-condensed phase is bound by the aptamer (considering an aptamer–FUS affinity of *K*_d_ *=* 280 ± 20 nM as determined above), and assuming the protein in the condensate phase is in equilibrium with the aptamer binding reaction, the protein concentration in the condensate phase amounts to *c*_FUS, condensate_ = *c*_FUS, total_ – *c*_FUS, monomer_ = 1414.6 ± 101.0 nM. From our volume fraction calculation, we estimated that the mean volume fraction of the condensates is *^ϕ^*^̄^ condensates= 1.55%. From these values, we determined the concentration of FUS in the condensates as *c*_FUS, condensate_ = 90.9 ± 0.7 μM. Multiplied by the molecular weight of the FUS-SNAP protein used (94.4 kDa), we obtain a protein mass concentration of 8.6 ± 0.6 mg/mL, comparable to what is expected for fragile protein–RNA condensates,^68^ although notably less dense than homotypic FUS condensates alone.^69^ This lowering of density is similar to the significant reduction in FUS concentration in condensates upon the addition of RNA previous reported.^63^ Of note, by dividing the total concentration of FUS in the condensate by the RNA concentration, we further obtain the FUS/RNA ratio inside the condensate, *c*_FUS, condensate_/ *c*_aptamer, FUS condensate_ = 1414.6 nM / 3.5nM = 40, indicating that in these condensates there is a large excess of protein over RNA.

Overall, we have demonstrated here that the DigitISA platform can simultaneously provide information on the affinity of the monomer–aptamer interaction, determine the concentration of native protein condensates and their mass densities, detect the stochiometric ratio of aptamers per condensate, and determine the partitioning coefficient of aptamer into protein condensates as well as the protein/RNA ratio inside the condensate. Quantifying dual binding affinities between a probe and a protein monomer and higher-order assembly would not be addressable by conventional binding assay techniques such as ELISA simply because it would be impossible to distinguish being the two different bound species as they would both contribute to the same fluorescent signal. Only by counting molecules and assemblies on a single particle level, can binding affinities, concentrations, and partitioning coefficients in a probe–analyte assay be simultaneously quantified in a heterogeneous multicomponent sample, as we show through our use of DigitISA to study a FUS–aptamer interaction. We anticipate that this could have widespread future applications in sensing native condensates, and could be a useful tool for the search and optimization of affinity probes to protein condensates. Such an assay has huge therapeutic potential as protein condensates are increasingly gaining attraction as therapeutic targets in a range of diseases.^70^

## Discussion

By combining microchip electrophoretic separation with single-molecule detection, we have demonstrated here a surface-free sensing platform for the digital detection and quantification of protein targets in solution. The DigitISA platform operates entirely in solution, does not require washing steps, and performs protein detection with single-molecule sensitivity in a single step using only a single affinity reagent. The assay format further combines affinity selection with physical separation and thus provides an additional, orthogonal criterion for target detection to afford selectivity in the sensing process, which further opens up the possibility to discriminate probe binding to multiple different species, including non-specific binding. We note that the inclusion of an electrophoresis step is similar to the multiplicative capability of achieving selectivity in surface-based assays through the use of an affinity reagent pair. Yet, given that DigitISA uses only a single affinity reagent per target, the complexity of assay design is reduced because validated, non-cross-reactive affinity probe pairs and multi-epitopic targets are not required. The surface-free nature of the assay also reduces false-positive signaling by non-specific surface adsorption. Hence, DigitISA realizes an assay design with a number of qualitative differences over classical surface-based sensing designs (see Table 1 and Figure 8).

**Figure 8:**
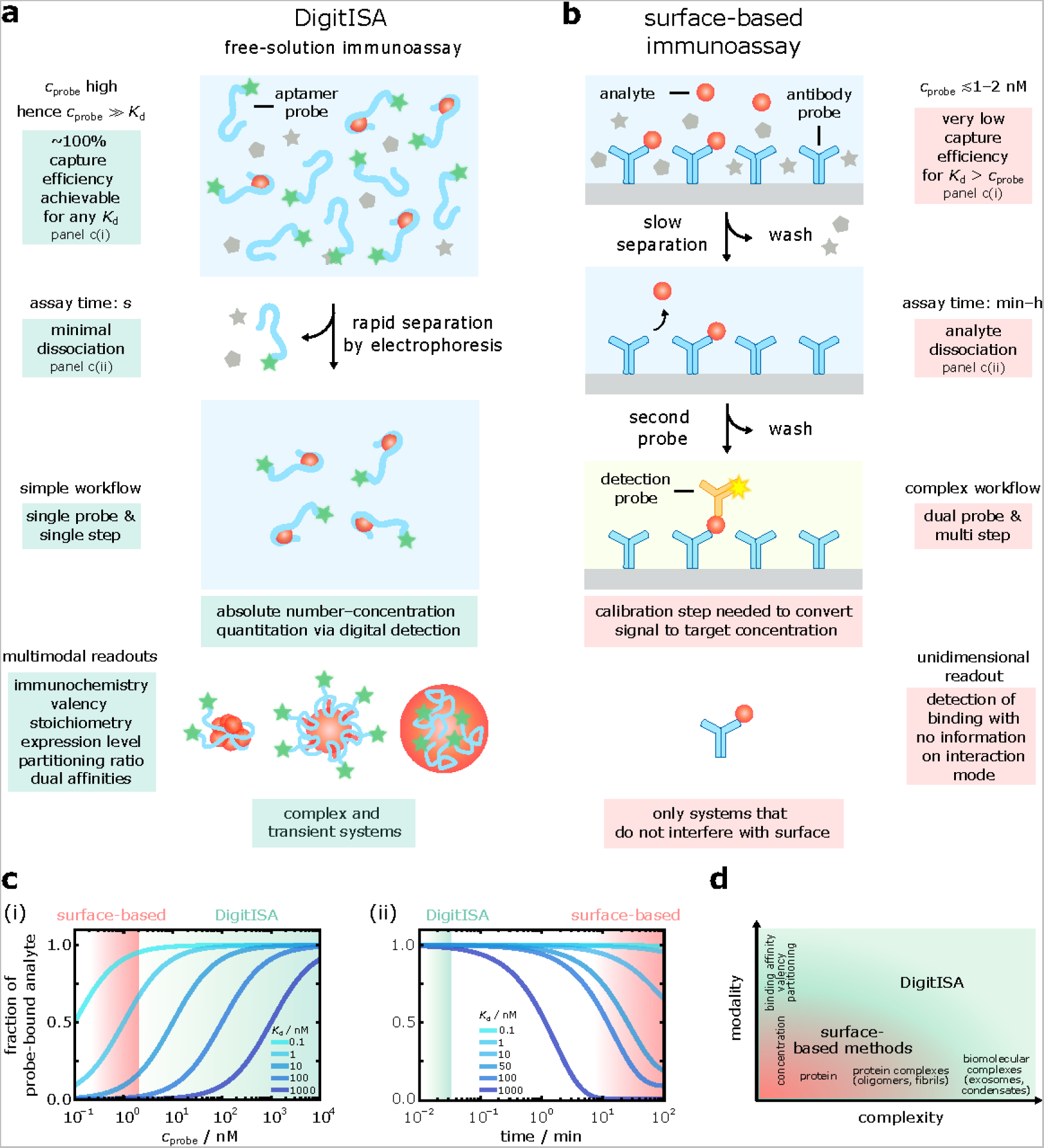
Comparison of the DigitISA assay with surface-based sensing methods. **(a)** Schematics depicting the principles of DigitISA and its advantages over surface-based methods. By performing the sensing reaction in solution, high concentrations of the affinity probe can be used, which permits quantitative target binding, even for probes with dissociation constants *K*_d_ > 1 nM (c(i)). Rapid removal of non-target bound probe by electrophoresis prevents the system from re-equilibrating (c(ii)) and sets the basis for quantitative analysis. The assay can be accomplished in a single step and requires only a single affinity reagent. Due to the digital nature of detection, absolute number–concentration quantification is possible, and additional parameters such as immunochemistry, stoichiometry, partitioning ratios and dual affinities can be extracted. DigitISA also allows for the analysis of complex and transient systems. **(b)** Schematic of surface-based immunosensor assays and their limitations. Conventional methods are limited to surface-capture probe concentrations (*c*_probe_) in the low nanomolar regime. Under these conditions, a significant amount of the analyte is not bound and thus remains undetected, especially when using affinity probes with *K*_d_ > 1 nM (c(i)). The binding equilibrium is disturbed during long washing steps. Hence, dissociation of the complex becomes significant, particularly for low affinity probes with *K*_d_ > 1 nM (c(ii)). Conventional assays involve multi-step procedures and typically require additional probes for detection and specificity. Detection modes require calibration to convert the signal to target concentration. The information content on target binding interaction is low and only targets that do not interfere with the surface are assayable. **(c)** Speciation curves depicting fraction of probe-bound analyte versus affinity probe concentration (c(i)) and fraction of probe-bound analyte versus time (c(ii)). Curves for probe–analyte affinities with *K*_d_ = 0.1–1000 nM (from light to dark blue) are shown. Green and red shaded areas denote operation regimes. **(d)** The multimodal detection capabilities of DigitISA augment the information content from sensing experiments beyond what is achievable and assayable with classical techniques. The nature of systems that can be studied are further expanded with DigitISA, enabling the study of transient systems such protein oligomers and liquid condensates.

**Table 1:** Comparison between the digital immunosensor assay (DigitISA) developed here and conventional surface-based immunosensor assays for protein detection.

| Digital immunosensor assay (DigitISA) | ELISA and bead-based immunoassays |
| --- | --- |
| Free solution | Surface-based |
| Complete binding of target molecules achievable even with low affinity probes | Capture efficiency determined by $K_d$ and surface concentration of affinity probe |
| Single-step assay with fast removal of excess affinity probe | Multi-step assay involving washing steps for removal of excess probes and reagents |
| Specificity from combined affinity and electrophoretic criteria | Specificity from affinity criteria only |
| Single affinity reagent | Affinity reagent pair |
| Absolute number–concentration quantitation through digital detection | Generally reliant on calibration to convert signal to target concentration |
| Multimodal readouts with information on immunochemistry/valency and stoichiometry, expression level etc. | Simple binding scenarios and limited information on binding modes |
| Complex and transient systems such as oligomers and liquid condensates | Only targets that do not interfere with surface |

One essential aspect of the DigitISA platform, and its assay design, is that it releases fundamental constraints of conventional immunosensing approaches in terms of the thermodynamics and kinetics of the immunoprobe–analyte interaction (Figure 8a–c, see also Supplementary Note 1). Due to the in-solution nature of the assay, near-quantitative binding of the target molecule can be achieved through the use of high probe concentrations, which together with the fast timescale of excess probe removal (∼2 s), enables the probe–analyte binding interaction to be maintained during the entire sensing process. Crucially, these two features combined allow quantitative protein biomarker sensing even with low affinity and rapidly dissociating capture probes such as aptamers in quantitative single-molecule sensing, as demonstrated throughout this work. This finding is significant as our approach allows quantitative protein sensing even at concentrations far below the *K*_d_ of the probe–analyte interaction. We note that aptamer probe concentrations in the picomolar to nanomolar regime were sufficiently high in our experiments to allow quantitative detection of the probe–target complexes under scrutiny; however higher concentrations would become necessary and could be used if even higher *K*_d_ value (i.e., lower affinity) probes were employed. In a previous work, we have demonstrated excess probe binding in a bulk assay format at micromolar probe concentrations, however, without the benefits of single-molecule detection.^71^

The implementation of the DigitISA approach in this work is based on aptamers as affinity probes. Even though antibodies are typically used as gold standard affinity reagents in conventional protein sensing applications, aptamers offer a number of advantages over antibodies, which are of particular relevance for the implementation of our DigitISA approach. Firstly, aptamers are favorable due to their higher electrophoretic mobility compared to antibodies, which facilitates separation between probe bound and unbound species. Secondly, aptamers offer recognition capabilities that rival those of antibodies, which is important to increase specificity because of the single probe assay format. And thirdly, the production of aptamers through in vitro evolution^29–31^ and chemical synthesis is fast and inexpensive, as compared to the often challenging production of antibodies. Finally, we would like to note that to date aptamers have found limited use in sensing applications mainly due to their relatively weak binding affinity and relatively fast rate of probe–analyte dissociation. The DigitISA assay, due to its design principle, overcomes this challenge and permits the use of these often-overlooked affinity reagents, because high concentrations of the aptamer can be used in the assay due to the in-solution nature of the DigitISA assay and the rapid separation of excess probe, as detailed above. Nonetheless, to complement the aptamer-based detection of targets, in future designs, it would be interesting to develop antibody-based probes for DigitISA experiments.

DigitISA provides additional qualitative advantages due to its digital detection modality, the multimodal information that can be obtained from the experiments, and the nature of systems that can be studied. While many protein-quantification assays, including conventional ELISAs, rely on calibration to convert a signal to a target concentration, the platform developed here counts the number of passing molecules in a digital manner and therefore enables absolute number–concentration quantitation. Specifically, we have shown for two examples, namely, the biotin–streptavidin complex and the IgE–aptamer complex that the assay is directly quantitative by recovering the nominal starting concentration, thereby validating our digital counting approach for the concentrations tested. In addition to direct readouts of absolute concentrations, DigitISA also enables characterization of immunochemistry valency and stoichiometry in a single experiment, as exemplified with the sensing of α-synuclein oligomers. Furthermore, our data on exosome biomarker detection shows that the DigitISA platform is well suited for the analysis of proteins in complex sensing scenarios. We simultaneously quantified the concentration of exosomes along with the expression levels of CD63, features that are hard if not impossible to obtain by other conventional methods. Moreover, we have shown that transient systems such as liquid condensates, which are also of increasing interest from a therapeutic point of view and fundamentally not addressable by classical protein sensing assays, are amenable to detailed analysis using our DigitISA assay. We have demonstrated that the platform can detect condensate concentrations and simultaneously provide information on the affinity of the monomer–aptamer interaction, determine the concentration of native protein within condensates and their mass densities, detect the stochiometric ratio of aptamers per condensate, and determine the partitioning coefficient of aptamer into protein condensates. Together, the multimodal detection capabilities of the DigitISA assay augment the information content from protein sensing experiment beyond what is achievable and assayable with classical techniques (Figure 8d). We note that the analytes tested in this work here were probed in aqueous buffer solutions. However, the DigitISA assay should be adaptable to the detection of targets in biological and clinical samples. In a previous work,^71^ we have demonstrated that an electrophoresis-based assay design is robust enough to function in complex biological media, with target selectivity demonstrated in cell lysate. Hence, we anticipate that the DigitISA will perform effectively in complex biological media including clinical samples where minimal sample pre-treatment is required.

A key aspect of the DigitISA technique is that a sufficient change in the charge-to-mass ratio (i.e., electrophoretic mobility) is required upon binding of the affinity probe to the target protein to allow for effective separation. We have previously shown that with the chip design used in our assay, a difference in mobility of around 0.5 · 10^−8^ m^2^ V^−1^ s^−1^ is sufficient to ensure separation.^72^ For the typical sizes of protein targets (hydrodynamic radius of around 3–4 nm), this mobility value corresponds to an absolute charge difference of about 2–3 units – a charge shift that is exceeded when a negatively charged aptamer of tens of nucleotides (each of which carries one negative charge per nucleic acid residue) is bound to a protein target. Hence, the relatively small physical size and high charge density of aptamers lead to a sufficient alteration of the electrophoretic mobility upon binding to a protein target and thus allows effective separation of the protein bound and non-bound forms. While our assay achieves good separation, improvements such as controlled injection^20^ or the use of a device design where flow and applied field are parallel to each other (e.g., via on-chip capillary electrophoresis)^73, 74^ are expected to provide some means to enhance the separation power further.

In terms of sensitivity, our DigitISA platform achieves quantitation with picomolar sensitivity comparable to that of ‘gold-standard’ ELISAs.^4^ Exemplary estimates for the limit of detection of the DigitISA approach are given in Supplementary Note 2. We note that more complex implementations of traditional assays, such as digital ELISAs can go below this sensitivity limit (i.e., sub-picomolar concentrations), an application to which the single-molecule-counting ability of DigitISA could be well suited. We predict that the sensitivity of DigitISA can be improved, for example, by increasing the integration time in data acquisition or by optimizing the window of detection (see Supplementary Note 3). According to Poisson sampling statistics, longer acquisition times decrease the magnitude of the standard deviation of the sample relative to the mean and thus reduce detection limits of the assay, providing a promising route towards facile protein sensing with femtomolar sensitivity. Similar arguments hold true for optimizing the detection window into regions where the signal-to-noise ratio is increased and the standard deviation in control sample is minimized. Furthermore, we would like to note that the detection range of the DigitISA assay is not limited to the single-molecule regime. In a previous work, we have demonstrated excess probe binding in a bulk assay format at micromolar probe concentrations,^40^ however, without the benefits of single molecule detection. Thus, the method can sense in the range from picomolar to micromolar concentrations.

The implementation of the DigitISA platform as demonstrated here is based on the integration of microfluidic free-flow electrophoresis with in-situ confocal microscopy. Future iterations of the platform could also incorporate, for example, multicolor single-molecule spectroscopy and FRET techniques,^75–77^ as well as other microfluidic separation modalities^73, 74^ combined with downstream analyses,^78^ to enhance separation power, assay parallelization, sensitivity, specificity, and robustness to experimental noise. We also foresee that DigitISA has the potential to be developed into a commercial, benchtop instrument, similar to those developed, for example, by Quanterix, Nanostring, or Fluidic Analytics. Given DigitISA’s key features, in particular that it allows probing a much broader range of interactions than surface-based immunoassays and goes beyond monovalent binding interactions, we anticipate that the DigitISA technology will have a demand in biomolecular analysis and diagnostics as an orthogonal tool to surface-based assays in the many situations where they do not work well.

Despite the advances of the DigitISA platform, certain fundamental limitations still exist. Like other techniques, including ELISA-like approaches, DigitISA requires a probe and is thus limited in performing explorative hypothesis-free studies. Moreover, DigitISA in its current implementation is not high throughput, however, engineering approaches are available that can transform chip-based techniques into higher throughput techniques, as is the case, for example, in surface plasmon resonance. Also, currently orthogonal knowledge of the *K*_d_ value is required for an experiment to not require *K*_d_ calibration. Finally, before widespread applicability with a dedicated instrument, DigitISA requires expertise in advanced microfluidics and single-molecule optics for its implementation, and the provision of equipment. However, we would like to note that ELISAs and other surface-based methods typically also require the acquisition of a plate reader or other optical instruments, which also means a significant investment.

Taken together, the microchip DigitISA platform presented herein constitutes a new experimental paradigm for protein biomarker sensing. We have demonstrated that the method is at least on par in terms of sensitivity with more traditional assay (i.e., in the pM range), yet with a number of qualitative advances. DigitISA not only enables highly sensitive number–concentration quantitation of proteins in complex sensing scenarios, but also enables characterization of immunochemistry valency and stoichiometry, expression levels, as well as detection of multiple protein states, determination of partitioning coefficient, and dual binding affinities, in a single measurement and in a digital manner. We also demonstrated the application to a liquid protein system that would be inherently not addressable by conventional sensing assays. We anticipate that the DigitISA methodology has the potential to find broad applicability in the detection and quantification of biomolecular and biomedically relevant targets, thus paving the way for DigitISA to become an orthogonal tool for sensitive multimodal biomolecular analysis and diagnostics, especially in the many situations where more traditional assays do not work well.

## Methods

### DigitISA platform

The DigitISA optofluidic platform integrates microchip free-flow electrophoresis with single-molecule confocal fluorescence microscopy. Schematics of the microfluidic separation device, the optical setup, and their integration are shown in Figure 2. Briefly, the electrophoresis device is fabricated in PDMS by standard soft-lithography and molding techniques and bonded to a glass coverslip (see below). Fluids are injected into the chip via polyethylene tubing from glass syringes (Hamilton) and solution flow rates are controlled by automated syringe pumps (Cetoni neMESYS). Fluid waste is guided out of the device by tubing inserted into device outlets. Importantly, the free-flow electrophoresis functionality of the device is attained by a design that makes use of liquid electrolyte electrodes,^19, 24^ where an electric potential is applied outside and downstream of the microfluidic separation chamber in separate electrolyte channels, and thereby allowed to propagate back to the separation area through the use of a co-flowing, highly conductive electrolyte solution (i.e., 3 M KCl). This capacity is realized in the chip architecture by connecting the main electrophoresis chamber, which harbors the sample and the flanking buffered carrier medium from both sides, with the co-following electrolyte channels via narrow bridges. These connectors control the transfer of the electrolyte to the electrophoresis chamber and allow a thin sheet of electrolyte to form at and flow along both edges of the chamber, acting as liquid electrodes. Electric potentials are applied from a power supply (EA-PS 9500-06, EA Elektro-Automatik) via hollow metal dispensing tips (20G, Intertonics) inserted into the electrolyte outlets and carried into the device via the electrolyte solution, resulting in an electric field that spans across the main separation channel. Crucially, the use of liquid electrodes allows any generated electrolysis products to be flushed out of the chip and Joule heating to be reduced, permitting strong electric fields to be applied in a stable manner. This feature is of utmost importance for effective and high-performance electrophoretic separation.^19, 20, 24^ Extensive characterization of the microfluidic chip was done in Saar et al.,^19^ and the chip architecture has since been used in a variety of studies.^20, 24, 25, 72, 79, 80^ Calibration of the device resistivity demonstrated a typical voltage efficiency of 30–40%.^19, 24, 25^ A variant of the microfluidic free-flow electrophoresis device was utilized for the oligomer and condensate DigitISA experiments. This device varied from the standard device by the addition of a desalting module, upstream of the electrophoresis chamber, in order to rapidly decrease the salt concentration on chip prior to electrophoretic analysis.^40^ By diffusional mixing of the sample with water, a 15 times reduction in salt concentration in the sample solution can be achieved, while 70% of the large samples (radius > 10 nm) are able to remain in the central sample flow and finally enter the electrophoresis chamber.^40^

The optical unit of the DigitISA platform is based on a laser-induced fluorescence confocal microscope optimized for microfluidic experiments. The electrophoresis chip is secured to a motorized scanning stage (PZ-2000FT, Applied Scientific Instrumentation (ASI)), which is mounted onto a ‘rapid automated modular microscope’ (RAMM) frame (ASI). The motorized *x*,*y*,*z*-stage is equipped with a *z*-piezo for controlling precise sample placement along the optical axis of the microscope. To excite the sample in the device, the beam of a 488-nm wavelength laser (Cobolt 06-MLD, 200 mW diode laser, Cobolt) is passed through a single-mode optical fiber (P3-488PM-FC-1, Thorlabs) and collimated at the exit of the fiber by an achromatic collimator (60FC-L-4-M100S-26, Schäfter + Kirchhoff) to form a beam with a Gaussian profile. The beam is then directed into the microscope, reflected by a dichroic beamsplitter (Di03-R488/561, Semrock), and subsequently focused to a concentric diffraction-limited spot in the microfluidic channel through a 60x-magnification water-immersion objective (CFI Plan Apochromat WI 60x, NA 1.2, Nikon). The emitted light from the sample is collected via the same objective, passed through the dichroic beam splitter, and focused by achromatic lenses through a 30-µm pinhole (Thorlabs) to remove any out-of-focus light. The emitted photons are filtered through a band-pass filter (FF01-520/35-25, Semrock) and then focused onto a single-photon counting avalanche photodiode (APD, SPCM-14, PerkinElmer Optoelectronics), which is connected to a TimeHarp260 time-correlated single photon counting unit (PicoQuant).

### Fabrication of the electrophoretic devices

The microfluidic device was designed using AutoCAD software (Autodesk) and printed on acetate transparencies (Micro Lithography Services). The replica mold for fabricating the device was prepared through a single, standard soft-lithography step^81, 82^ by spinning SU-8 3025 photoresist (MicroChem) onto a polished silicon wafer to a height of around 25 µm. The UV exposure step was performed with a custom-built LED-based apparatus^83^ and the precise height of the features were measured to be 28 µm by a profilometer (Dektak, Bruker). The mold was then used to generate a patterned PDMS slab. To this effect, the mold was casted in a 10:1 (*w*/*w*) mixture of PDMS (Dow Corning) and curing agent (Sylgard 184, Dow Corning), degassed and baked for 1.5 h at 65°C. The formed PDMS slab was cut and peeled off the master and access holes for the inlet tubes were introduced using biopsy punches. The devices were then bonded to a thin glass coverslip after both the PDMS and the glass surface had been activated through oxygen plasma (Diener electronic, 40 % power for 15 s). Before injecting the solutions into the channels, the chips were exposed to an additional plasma oxidation step (80 % power for 500 s) which rendered the channel surfaces more hydrophilic.^84^

### Protein preparation

The monovalent streptavidin sample was kindly provided by the Howarth Lab (University of Oxford),^23^ and recombinant IgE Kappa (clone AbD18705_IgE, Cat#HCA190) was purchased from Bio-Rad Laboratories.

Wild-type α-synuclein was recombinantly expressed and purified according to published procedures.^85, 86^ Briefly, *Escherichia coli* BL21(DE3) cells were transformed with a pT7-7 expression vector encoding the gene for human wild-type α-synuclein. Cultured cells were grown at 37°C under shaking in Luria Bertani medium supplemented with Ampicillin (100 μg/mL) to an optical density at 600 nm of 0.6. Expression was induced with 1 mM isopropyl β-D-thiogalactoside (IPTG). After over-night incubation at 28°C under shaking, cells were collected by centrifugation, washed in phosphate buffered saline (PBS) followed by a further round of centrifugation, resuspended in lysis buffer (10 mM Tris-HCl (pH 7.7), 1 mM EDTA, protease inhibitor), and lysed by sonication on ice. Following centrifugation, heat-sensitive proteins were precipitated out of the lysate supernatant by boiling (20 min, 80–95°C), and subsequently removed by centrifugation. DNA was precipitated out by incubation with streptomycin sulphate (10 mg/mL, 15 min, 4 °C), and removed by centrifugation. α-synuclein was precipitated out of the supernatant by the slow addition of ammonium sulphate (361 mg/mL) while stirring (30 min, 4°C). The pellet containing α-synuclein was collected by centrifugation and resuspended in 25 mM Tris-HCl (pH 7.7) buffer. Following dialysis using a 3500 MWCO dialysis membrane (Thermo Fisher) against 25 mM Tris-HCl (pH 7.7) buffer at 4°C, the protein was subjected to ion exchange chromatography and loaded onto a HiLoad 26/10 Q Sepharose high performance column (GE Healthcare), and eluted at ∼350 mM NaCl with a salt gradient from 0 M to 1.5 M NaCl in buffers containing 25 mM Tris-HCl (pH 7.7). Selected fractions were subsequently subjected to size exclusion chromatography (SEC) using a Superdex 75 26/60 column (GE Healthcare) and eluted in PBS (pH 7.4) buffer. All chromatography steps were done at 20°C. Protein concentration was determined by absorbance at 275 nm, using an extinction coefficient of 5600 L/mol·cm. Protein sample was aliquoted, flash-frozen, and stored at –80°C.

Expression and purification of FUS protein was adapted from Patel et al.^64^ Briefly, recombinant FUS was expressed in Sf9 insect cells (Expression Systems, Cat#94-001F) for 72 h using the baculovirus expression system^87^ and produced as a C-terminal SNAP fusion with an N-terminal maltose-binding protein (MBP) and hexahistidine (His6) tags. Cells expressing His6-MBP-FUS-SNAP were lysed using the LM10 Microfluidizer Processor (Analytik) in lysis buffer (50 mM Tris-HCl (pH 7.4), 500 mM KCl, 5% (w/v) glycerol, 10 mM imidazole, 1 mM DTT, 12.5 ng/ml benzonase), supplemented with EDTA-free protease inhibitor cocktail (Roche Applied Science). The soluble fraction was collected after centrifugation for 45 min at 4°C. The soluble fraction was incubated with Ni-NTA resin (Qiagen) for 1 h, and then washed with 10 column volumes wash buffer (50 mM Tris-HCl (pH 7.4), 500 mM KCl, 5% (w/v) glycerol, 10 mM imidazole, 1 mM DTT). The protein was eluted with an imidazole buffer (50 mM Tris-HCl (pH 7.4), 1 M KCl, 5% (w/v) glycerol, 300 mM imidazole, 1 mM DTT), and then incubated with amylose resin (New England Biolabs) for 1 h. The resin was washed with 10 column volumes imidazole buffer and then eluted with maltose buffer (50 mM Tris-HCl (pH 7.4), 1 M KCl, 5% (w/v) glycerol, 500 mM arginine, 50 mM maltose, 1 mM DTT). The eluted sample was concentrated with Amicon Ultra centrifugal filters (Merck Millipore). The His6 and MBP tags were cleaved using the 3C preScission protease (provided by the Max Planck Institute of Molecular Cell Biology and Genetics (MPI-CBG) protein expression facility) through 4 h incubation at room temperature. Sample was then applied to SEC using a HiLoad 16/600 Superdex 200 pg (GE Life Sciences) on an Akta Pure (GE Life Sciences) system in storage buffer (50 mM HEPES, 1 M KCl, 5% (w/v) glycerol, 1 mM DTT) and the fractions containing FUS-SNAP were pooled. Sample was then concentrated with Amicon Ultra centrifugal filters (Merck Millipore), flash-frozen, and stored at –80°C.

### Exosome preparation

MDA-MB-231 epithelial breast cancer cells (ATCC, Cat#HTB-26) were cultured according to supplier’s instructions with Dulbecco’s Modified Eagle Medium (DMEM; Thermo Fisher) and supplemented with 10% (v/v) fetal bovine serum (Merck), 50 U/mL penicillin, 50 μg/mL streptomycin, and 1% (v/v) GlutaMAX (Thermo Fisher). Cells were incubated at 37°C with 5% CO2. Cells were cultured until 80% confluent before passaging and did not exceed 15 passages. For exosome harvesting, MDA-MB-231 cells were cultured until 80% confluent, washed twice with PBS and incubated with serum-free culture medium for 48 h before the conditioned medium was collected for further processing. Cell were mycoplasma tested every 6 months.

To isolate exosomes for DigitISA experiments, conditioned medium from MDA-MB-231 cells was processed using an adapted protocol as described.^88^ In brief, conditioned medium was centrifuged at 4°C for 30 min to remove cells and cellular debris, followed by filtration through a syringe-operated 0.2 μm filter unit (Millipore) to remove non-exosomal extracellular vesicles. The filtrate was centrifuged at 4 °C for 4 h to pellet out exosomes. The pellet was resuspended in 500 μL 10 mM HEPES and stored at 4 °C pending analysis. Exosome samples were used within 5–7 days of isolation.

To characterize exosomes by transmission electron microscopy (TEM), exosome samples were visualized by TEM using negative staining method. 10 μL sample was applied on glow-discharged carbon-coated copper grids for 2 min, washed twice in ultrapure water for 30 s each, and stained with 1% (w/v) water solution of uranyl acetate for 2 min at room temperature. Imaging was performed on a FEI Tecnai G20 electron microscope, operating at 200 kV and with a 20 µm objective aperture, and images were recorded with an AMT camera.

For extraction of proteins from exosomes for Western blotting, cultured cells were washed twice with cold PBS and solubilized in 1 mL radio immunoprecipitation assay buffer (RIPA; Thermo Fisher). Lysis and extraction buffer were supplemented with 10 μL Halt Protease Inhibitor Cocktail and 10 μL EDTA (Thermo Fisher) immediately before use. Lysed samples were incubated on ice for 5 min, collected into a microcentrifuge tube and centrifuged at 4°C for 15 min to pellet cell debris. The supernatant (cell lysate) was collected and stored at –80°C pending further analysis. Exosome protein samples were prepared by resuspending pellets in the supplemented RIPA buffer.

For Western blotting, isolated exosome samples) were lysed in lithium dodecyl sulfate (LDS) sample buffer (4X Bolt; Invitrogen), denatured by heating for 5 min at 95°C and subjected to electrophoresis using precast NuPAGE Novex 4–12% Bis-Tris Proteins Gels (Invitrogen) in NuPAGE MES SDS Running Buffer (Invitrogen) under non-reducing conditions at 200 V for 45 min. Proteins were electrotransferred onto PVDF Transfer Membranes (Thermo Fisher) using NuPAGE Transfer Buffer (Invitrogen). Membranes were blocked with 5% (w/v) bovine serum albumin in phosphate-buffered saline (Thermo Fisher, Oxoid) with 0.1% (v/v) Tween-20 (PBST) for 1 h. Membranes were subsequently probed with mouse anti-CD9 (Ts9) (Invitrogen, Cat#10626D, 1:1000), mouse anti-CD63 (Ts63) (Thermo Fisher, Cat#10628D, 1:1000) and rabbit anti-calreticulin [EPR3924] (Abcam, Cat#ab92516, 1:1000) monoclonal primary antibodies for 1 h in PBST followed by incubation with HRP-conjugated anti-mouse IgG H&L (Invitrogen, Cat#A16078, 1:1000) and anti-rabbit IgG H&L (Abcam, Cat#ab7090, 1:1000) secondary antibodies in PBST for 1 h. All incubations were carried out on a rotating wheel drum at room temperature. Western blots were washed three times in PBST for 5 min after each incubation step and were visualized using Pierce enhanced chemiluminescence Western blotting substrate (Thermo Fisher) on a G:BOX Chemi XX6 gel documentation system (Syngene).

### Sample preparation for DigitISA experiments

The complex between the biotinylated and fluorophore-conjugated DNA strand (5’-Atto488-CGACATCTAACCTAGCTCACTGAC-Biotin-3’, HLPC purified; Biomers) and monovalent streptavidin was formed by mixing 25 pM of the monovalent streptavidin sample with 50 pM of biotinylated probe DNA in 10 mM HEPES (pH 7.4) buffer (Sigma) supplemented with 0.05 % Tween-20 (Thermo Scientific). Prior to its injection to the chip, the mixture was incubated at room temperature for 5 minutes.

Recombinant IgE Kappa was dissolved to a concentration of 40 nM in 10 mM HEPES (pH 7.4) buffer supplemented with 0.05 % (v/v) Tween-20. The protein sample was then mixed with the aptamer probe (5’-Atto488-TGGGGCACGTTTATCCGTCCCTCCTAGTGGCGTGCCCC-3’, HPLC purified; Integrated DNA Technologies (IDT)) diluted to a concentration of 50 pM and incubated for 10 minutes before the experiment at room temperature prior to their injection to the chip.

α-synuclein fibrils were detected using T-SO508 aptamer^36^ (5’-Alexa488-TTTTGCCTGTGGTGTTGGGGCGGGTGCG-3’, HPLC purified; IDT). Prior to its use, the aptamer (100 µM stock in 1X TE buffer) was heated to 70°C and cooled to room temperature to facilitate correct folding. The α-synuclein fibrils were prepared following previously published procedures.^42^ Briefly, α-synuclein monomer was incubated at 70 μM in PBS (pH 7.4, 0.1 M ionic strength) at 37 °C under constant agitation for 4–6 days. The sample was then centrifuged and the fibrillar pellet washed twice with PBS. Following sonication (10 % power, 30 % cycles for 1 min; Sonopuls HD 2070, Bandelin), the fibrils were spun down and re-suspended in 10 mM HEPES (pH 7.4), 0.05 % (v/v) Tween-20. The aptamer and the fibrils were then mixed by suspending them into 10 mM HEPES (pH 7.4) buffer supplemented with 0.05 % (v/v) Tween-20 to final concentrations of 10 nM and 80 nM (monomer equivalent), respectively. The mixture was incubated for 10 minutes before its injection to the chip.

α-synuclein oligomers were detected using 100nM of T-SO606 aptamer^36^ (5’-Alexa488-TTTTGGGTCGGCTGTCCGTGGGTGGGGA-3’, HPLC purified; IDT). Prior to its use, the aptamer (100 µM stock in 1X TE buffer) was heated to 70°C and cooled to room temperature to facilitate correct folding. Oligomers were formed by incubating wild-type α-synuclein (100 μM) in PBS buffer (pH 7.4), at 37 °C with shaking at 300 rpm over 12 h, following published procedures.^40^ An aliquot was withdrawn and centrifuged for 15 min at 21’000 *g*, to pellet insoluble, fibrillar components of the reaction mixture. The supernatant, containing monomeric and oligomeric α-synuclein, was carefully removed and used immediately for DigitISA experiments.

Exosomes isolated from MDA-MB-231 cells were detected using a CD63-specific aptamer (5’-CACCCCACCTCGCTCCCGTGACACTAATGCTA-3’; HPLC purified, Biomers). Prior to its use, the aptamer (100 µM stock in 1X TE buffer) was heated to 70°C and cooled to room temperature to facilitate correct folding. The aptamer and the isolated exosomes were then mixed by adding them into 10 mM HEPES (pH 7.4) buffer supplemented with 0.05 % (v/v) Tween-20 to final concentrations of 5, 10 or 100 nM aptamer. The exosomes were diluted 2x from the stock. The mixture was incubated for 10 minutes before its injection to the chip.

FUS condensates were detected using the ‘GGUG’ RNA aptamer^66^ (5’-UUGUAUUUUGAGCUAGUUUGGUGAC-3’-Atto488, Biomers). Condensates were formed by diluting the protein to a final concentration of 2 μM in 50 mM TRIS-HCl (pH 7.4) at 25 mM KCl with 0.02% (v/v) Tween-20. After phase separation, the sample was mixed with 700 nM of aptamer and then allowed to equilibrate for 15 minutes before being injected into the chip. The sample was subjected to desalting on chip.^40^

### Experimental procedures

For experiments performed on the biotin–streptavidin system, the sample and the co-flowing buffer were injected into the microfluidic device at a flow rate of 70 and 2000 µL h^−1^, respectively, and the 3 M KCl electrolyte solution from each of its inlets at 300 µL h^−1^ using glass syringes. For DigitISA experiments on IgE and on the α-synuclein fibrils, these injection flow rates were 100, 1200, 400 µL h^−1^ and 50, 1200, 200 µL h^−1^, respectively. For DigitISA experiments on exosomes, the injection flow rates were 10, 1000, 300 µL h^−1^, respectively. α-synuclein oligomer and FUS condensate DigitISA experiments were run on the desalting variant of the FFE device. In this variant, the sample was injected into the device and surrounded by an additional desalting buffer, the sample then entered the central channel and was further surrounded by the co-flow buffer, as previously reported.^40^ For the condensates and oligomer, the flow rates were 30, 150, 800, 200 µL h^−1^, 10, 140, 1000, 200 µL h^−1^, respectively. The PDMS–glass chip was secured to the motorized, programmable microscope stage and once a stable flow in the device had been established, a potential difference across the device was applied. The photon-count time traces were obtained by translocating the microscope stage across the cross-section (Figure 2, step-scan line is indicated) using a custom-written Python script that simultaneously controlled the stage movement and the data acquisition at a distance of 4 mm downstream from where the electric field was first applied and at mid-height of the device (i.e., ∼14 µm above the surface of the glass coverslip). The direction of scanning was varied between Figs. 2–4 and 5–7 (note the direction of channel positions), but importantly the direction of electrophoresis was the same for all experiments. Each of the experiments was performed in a freshly fabricated PDMS device and simultaneous current readings were taken to ensure that the efficiencies did not vary between the devices and comparisons could be drawn between the deflected distances at which molecules eluted. The laser power at the back aperture of the objective was adjusted to 150 µW in all experiments, except for experiments on oligomers were a laser power of 370 µW was used, and experiments on condensates/exosomes were laser powers of 100 µW were used. We note that these laser powers ensure that the fluorophores (Alexa488, Atto488) are nearly saturated; hence, they emit the maximum number of photons per excitation cycle. This ensures that single-molecule events have sufficiently high signal-to-noise ratio to discriminate between photons that originate from single fluorescent molecules and those that correspond to background. Photon recordings were done in T2 mode and the arrival times of photons were measured in respect to the overall measurement start with 16-ps resolution.

### Data analysis

A burst-search algorithm was used to extract the number of molecules traversing the confocal volume from the recorded photon-count time trace using a combined inter-photon time (IPT) and photon-count threshold. The custom code (written in Python, version 3.7) is available as Supplementary Software or on the GitHub repository: https://github.com/rj380cam/digitISA. Briefly, the passing molecules were identified and distinguished from the background by requiring the IPT to remain short for the arrival of a number of consecutive photons. This approach has been shown to allow effective discrimination between background photons and those that originate from a fluorescently labeled molecules passing the laser spot and emitting photons.^21, 22, 89, 90^ Specifically, an IPT threshold of 100 µs was used for the analysis of the biotin– streptavidin interaction and the IgE sample with consecutive photon arrival events identified as a molecule when a packet of at least 7 photons arrived each with an IPT below threshold. For the analysis of the fibril, oligomer, and exosome samples these thresholds were set to 5 µs and 30 photons, respectively. In all cases, before analysis the IPT traces were processed with Lee filter (*n* = 4)^91^ to smoothen regions of constant signal while keeping those with rapid parameter changes, such as the edges of the bursts unaffected.

In the FUS condensate experiments, due the bulk signal stemming from the monomeric FUS– aptamer interaction, and the large burst signals from the detected condensates, a modified analysis method was carried out. In this modified method, the photon counts were binned into 1 ms intervals and each 5 s trace from a specific coordinate in the channel was analyzed by a thresholding algorithm. Burst were classified as regions where the signal exceeded a threshold unique to each trace (see the dashed line in Figure 7d). This threshold was 5 standard deviations above the mean (both parameters taken from the distribution of intensities for all 1 ms bins).

Graphing and plotting of data was done in Python (version 3.7) or Origin 2020 (OriginLab).

### Dimensions of the confocal volume from FCS

The dimensions of the confocal volume were determined by performing an FCS experiment on Atto488 carboxylic acid (100 pM; 150 µW laser power). Using its diffusion coefficient of *D* = 400 µm^2^ s^−1^, the effective volume of the confocal volume was evaluated to be *V*eff = 4.2 fL and the kappa factor to be *κ* = 6.0, yielding an estimate for the dimensions of the confocal volume as *z* = 3 µm in its height and around *w* = 0.4 µm in its width. The correlation analysis was done using the SymPhoTime 64 software package (Picoquant).

### Simulation of probe–target binding interaction

Estimations of analyte binding and complex dissociation depicted in Figure 8c were obtained by examining the binding of probe (P) to its target analyte (A) according to:

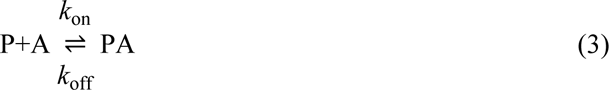

where *k*_on_ and *k*_off_ are the rate constants for the formation of the complex and its dissociation, respectively. The simulations in Figure 8c(i) describe the thermodynamic equilibrium for probes with various dissociation constants *K*_d_ ranging from 0.1 nM to 1000 nM as a function of its concentration *c*_probe_. The simulations in Figure 8c(ii) illustrate the dissociation of the protein– analyte complex after the removal of the excess reagents relative to the amount of complex that was present at equilibrium. Any changes in the dissociation constant (*K*_d_ = *k*_off_⁄*k*_on_) were assumed to originate from alterations in the rate constant that govern the dissociation of the complex 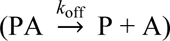 rather than changes in the rate constant governing the formation of the complex 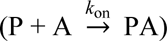. The rate constant for the latter reaction varies between probes.^92^ Here, we used *k*_on_ = 10^−4^ M^−1^s^−1^, which for a fixed *K*_d_ yields conservative estimates for *k*_off_, and hence, for the rate at which the complex dissociates.

## Statistics & Reproducibility

No statistical method was used to pre-determine sample size. No data were excluded from the analyses. The experiments were not randomized. The investigators were not blinded to allocation during experiments and outcome assessment.

## Data availability

All the data generated in this study are available within the main text and the Supplementary Information file. Source data are provided with this paper.

## Code Availability

Computer code used in this article for the analysis of photon time traces are available as Supplementary Software or on the GitHub repository: https://github.com/rj380cam/digitISA/

## Acknowledgements

The authors thank the Howarth Lab (University of Oxford) for monovalent streptavidin samples. The research leading to these results has received funding from the European Research Council under the European Union’s Horizon 2020 Framework Programme through the Marie Skłodowska-Curie grant MicroSPARK (agreement no. 841466; G.K.) and MicroProtLip (agreement no. 896068; M.A.C.), the Herchel Smith Fund of the University of Cambridge (G.K.), the Wolfson College Junior Research Fellowship (G.K.), the Schmidt Science Fellows program in partnership with the Rhodes Trust (K.L.S.), the Engineering and Physical Sciences Research Council (K.L.S), St. John’s College. (K.L.S), the EPSRC Cambridge NanoDTC (EP/L015978/1; W.E.A.), the European Union Horizon 2020 research and innovation programme (ETN grant 674979-NANOTRANS; Q.P.), the European Research Council under the European Union’s Seventh Framework Programme (FP7/2007-2013) through the ERC grants PhysProt (agreement no. 337969; T.P.J.K), and the Newman Foundation (T.P.J.K.). W.C.T. acknowledges funding from the Cambridge Commonwealth, European & International Trust at Cambridge University. T.J.W acknowledges funding from the Harding Distinguished Postgraduate Scholar Programme, administered by the Cambridge Commonwealth, European & International Trust at Cambridge University. The authors gratefully acknowledge the Cambridge Advanced Imaging Centre for their support & assistance in this work.

## Author contributions

G.K., K.L.S., W.E.A., and T.P.J.K. conceptualized the study and devised the methodology, G.K., K.L.S., P.C., C.G.T., and D.K. contributed experimental hardware and expertise, G.K., W.E.A., T.J.W., M.A.C., A.P. performed the investigation, G.K., K.L.S., W.E.A., T.J.W., R.P.B.J., and Q.P. performed the analysis, and R.P.B.J. and Q.P. contributed relevant software. W.C.T., A.K.J., L-M.v.d.L., T.M.F., and S.A. provided materials, expertise, and characterization data. G.K., K.L.S., W.E.A., and T.J.W. wrote the original draft and all other authors reviewed and edited it.

## Conflict of interest

G.K., K.L.S., W.E.A., and T.P.J.K. declare the following competing interests. Parts of this work have been the subject of a patent application filed by Cambridge Enterprise Limited, a fully owned subsidiary of the University of Cambridge. Inventors: Krainer, G.; Saar, K.L.; Arter, W.E., Knowles, T.P.J.; Applicant: Cambridge Enterprise Ltd.; Title: Highly sensitive biomolecule detection and quantification. Publication Number: WO/2021/176065; Publication Date: 10.09.2021; International Application No.: PCT/EP2021/055614; International Filing Date: 05.03.2021. The remaining authors declare no competing interests.

## Notes

### Summary of Updates

n/a

