## Supplementary Information for "Direct digital sensing of protein biomarkers in solution"

**Supplementary Note 1: Considerations on fundamental limitations of surface-based methods**

Protein sensing approaches that remain reliant on surface-immobilization of target analyte molecules via capture probes and multi-step washing protocols exhibit fundamental limitations due to the underlying thermodynamics and kinetics of the immunoprobe–analyte interaction. Principally, the concentration of affinity capture probes that can be utilized in such approaches is limited by the finite area of the surface. The maximally achievable surface-capture probe concentrations in surface-based methods is ~1–2 nM;[1,2] thus, for a capture probe with a dissociation constant ($K_d$) of 1 nM, as is typical of many affinity reagents, only 50 % of an analyte will be bound to the probe, which falls to <1 % for capture probes with $K_d$ > 100 nM (Figure 8c(i)). This low capture efficiency is exacerbated by dissociation of the probe–analyte complex (particularly for $K_d$ > 1 nM) as the binding equilibrium is disturbed over tens of minutes to hours by the washing steps required by the ELISA or bead-based protein detection assays; hence, off-rates become significant (Figure 8c(ii)). These lengthy assay workflows thus present challenges in maintaining probe–analyte interactions under non-equilibrium conditions. Together, these effects constitute intrinsic drawbacks to surface-based methods, resulting in the limitation that highly optimized capture probes with sub-nanomolar dissociation constants are required for effective sensing, and that a calibration step is needed that converts the recorded signal to the actual target concentration.

**Supplementary Note 2: Estimation of the limit of detection**

Due to the digital nature of the DigitISA assay and the possibility to obtain calibration-free readouts of molecular concentrations, the assay also allows for a direct estimation of the limit of detection (LOD). The LOD is defined as three standard deviations above the mean count of the control sample ($\text{LOD} = \bar{n}_{\text{control}} + 3 \cdot s_{\text{control}}$). For the example of monovalent binding of streptavidin to biotin (Figure 2, main text), with the control sample including

$\bar{n}_{\text{control}} = 789$ molecules that eluted in the detection region and with the standard deviation of the control sample being $s_{\text{control}} = 30$ counts, we estimate the LOD to be $789 + 3 \cdot 30 = 879$ molecules. Using the scaling ($\bar{n} = 62.8 \cdot c + 789$) of mean recorded counts ($\bar{n}$) versus molecular concentrations ($c$) calculated from the control sample ($\bar{n}_{\text{control}} = 789$, $c_{\text{control}} = 0$ pM) and the biotin–streptavidin sample ($\bar{n}_{\text{sample}} = 2391$, $c_{\text{sample}} = 25.5$ pM), we conclude the LOD to be 1.5 pM. By a similar argument, we estimated the LOD of the IgE–aptamer measurement (Figure 3, main text) to be 0.9 pM, and for the fibril sample (Figure 4, main text) to be 6.5 pM. This demonstrates that the DigitISA assay has sensitivities in the picomolar detection range. We note that the values we quote describe the LOD of the measurement system. The LOD of specific assays depends on the assay design and is in the picomolar range in the limit when high affinity probes are used and the majority of the target molecules are bound to the probe.

**Supplementary Note 3: Routes to improving the limit of detection**

We predict that the sensitivity of the DigitISA assay could be improved further by increasing the integration time in data acquisition, in particular by recording the passage of analytes at a single channel position where the signal-to-background ratio (*i.e.*, $\bar{n}_{\text{sample}}/\bar{n}_{\text{control}}$) is maximized and the standard deviation in the control signal ($s_{\text{control}}$) is minimized. According to the principles of Poisson sampling, for data acquisition time $t$, the mean counts scale with $\bar{n} \propto t$ whereas the standard deviation scales with $s \propto t^{1/2}$; hence, longer acquisition times will therefore decrease the magnitude of $s$ relative to $\bar{n}$ and reduce the LOD of the assay. For example, at the optimal channel position in the case of the monovalent–streptavidin experiment (Figure 2, $x = 1630$ μm, signal-to-background = 7.5, $s_{\text{control}} = 0.5$), we calculate that for 5 min acquisition time (the approximate time required for the step-wise scanning modality), such an approach would afford a LOD of 400 fM. Increasing the experiment time to 1 h, as is comparable with fast ELISA protocols, affords a predicted LOD of 50 fM. Thus, although accurate quantitation through this approach would require calibration of the observed signal against known standards, we envisage that the DigitISA platform provides a promising route towards facile protein sensing with sub-pM sensitivity. Moreover, sensitivity could be improved as well further by optimizing the region of data acquisition. For example, altering the acquisition window for the fibril-sensing experiment (Figure 4) to between $1600\ \mu m < x < 2100\ \mu m$ affords a ~10-fold reduction in LOD from 6.5 pM to 0.55 pM, although such an approach would require also comparison of the observed $\bar{n}_{\mathrm{control}}$ with calibration measurements to allow absolute quantification. We note that the detection limit is determined by the non-complete baseline separation of the bound and non-probe and, hence, LOD could be enhanced by further optimizing the resolution of the separation unit (see discussion in the main text).
